# Obesity promotes breast epithelium DNA damage in BRCA mutation carriers

**DOI:** 10.1101/2022.07.29.502090

**Authors:** Priya Bhardwaj, Neil M. Iyengar, Heba Zahid, Katharine M. Carter, Dong Jun Byun, Man Ho Choi, Qi Sun, Oleksandr Savenkov, Charalambia Louka, Catherine Liu, Phoebe Piloco, Monica Acosta, Rohan Bareja, Olivier Elemento, Miguel Foronda, Lukas E. Dow, Sofya Oshchepkova, Dilip D. Giri, Michael Pollak, Xi Kathy Zhou, Benjamin D. Hopkins, Ashley M. Laughney, Melissa K. Frey, Lora Hedrick Ellenson, Monica Morrow, Jason A. Spector, Lewis C. Cantley, Kristy A. Brown

## Abstract

Obesity is an established risk factor for breast cancer among women in the general population after menopause. Whether elevated bodyweight is a risk factor for women with a germline mutation in *BRCA1* or *BRCA2* is less clear due to inconsistent findings from epidemiological studies and lack of mechanistic studies in this population. Here, we show that DNA damage in normal breast epithelium of *BRCA* mutation carriers is positively correlated with body mass index and with biomarkers of metabolic dysfunction. Additionally, RNA-sequencing reveals significant obesity-associated alterations to the breast adipose microenvironment of *BRCA* mutation carriers, including activation of estrogen biosynthesis, which impacts neighboring breast epithelial cells. We found that blockade of estrogen biosynthesis or estrogen receptor activity decreases DNA damage, whereas treatment with leptin or insulin increases DNA damage in *BRCA* heterozygous epithelial cells. Furthermore, we show that increased adiposity is associated with mammary gland DNA damage and increased penetrance of mammary tumors in *Brca1*+/- mice. Overall, our results provide mechanistic evidence in support of a link between bodyweight and breast cancer development in *BRCA* mutation carriers and suggests that maintaining a healthy bodyweight or pharmacologically targeting estrogen or metabolic dysfunction may reduce the risk of breast cancer in this population.

**One Sentence Summary:** Elevated bodyweight is positively associated with DNA damage in breast epithelium of *BRCA* mutation carriers

## INTRODUCTION

Inheriting a pathogenic mutation in the DNA repair genes *BRCA1* or *BRCA2* is causally linked to the development of breast and ovarian cancer in women (*1, 2*). Although there is strong evidence linking obesity to the development of hormone receptor positive breast cancer after menopause in the general population (*3*), there are conflicting results in *BRCA* mutation carriers. Some studies have found that maintaining a healthy bodyweight or weight loss in young adulthood is associated with delayed onset of breast cancer (*4, 5*). Other studies have reported that adiposity or elevated bodyweight in adulthood is associated with increased cancer risk (*5–9*). Conversely, some reports indicate that increased body mass index (BMI) in young adulthood may have protective effects, and that risk is modified by menopausal status (*9–11*). The lack of clarity on the role of bodyweight and risk of breast cancer development in *BRCA* mutation carriers limits the ability of clinicians to provide evidence-based guidance on prevention strategies beyond prophylactic surgical intervention.

Weight gain and obesity are often coupled with metabolic syndrome, insulin resistance, and significant changes to adipose tissue, including that of the breast microenvironment (*12–15*). Obesity-induced changes to breast adipose tissue includes dysregulation of hormone and adipokine balance, and increased production of inflammatory mediators (*16*). For example, estrogen biosynthesis is increased in obese breast adipose tissue due to overexpression of aromatase in adipose stromal cells which catalyzes the conversion of androgens to estrogen (*17–19*). Additionally, excessive expansion of adipocytes leads to hypoxia, lipolysis, and altered adipokine production including higher leptin to adiponectin ratio (*15, 20, 21*). These changes to the breast microenvironment may have important implications for breast carcinogenesis given that breast epithelial cells are embedded in this milieu and engage in epithelium-adipose crosstalk (*22*).

BRCA1 and BRCA2 are critical for their role in homologous recombination-mediated repair of DNA double strand breaks (*23*). Mutations in either *BRCA1* or *BRCA2* cause a defect in DNA repair which can lead to an accumulation of DNA damage and consequently, tumorigenesis (*24, 25*). Studies have linked obesity or metabolic syndrome to DNA damage, including in leukocytes (*26*), skeletal muscle (*27*), peripheral blood mononuclear cells (*28*), and in pancreatic β-cells (*29*), but no studies have examined the relationship between obesity and DNA damage in normal breast epithelial cells.

We show that BMI and markers of metabolic dysfunction are positively correlated with DNA damage in normal breast epithelium of women carrying a *BRCA* mutation, a finding that is extended to the fallopian tube of *BRCA* mutation carriers. RNA-sequencing of whole breast tissue and of isolated breast epithelial organoids from *BRCA* mutation carriers, along with *ex vivo* and *in vitro* studies with *BRCA1* and *BRCA2* mutant primary tissues and cell lines, suggests several obesity-associated factors as possible drivers of DNA damage. Additionally, metformin, fulvestrant, leptin neutralizing antibodies and a PI3K inhibitor reduce damage induced by the obese breast microenvironment. *In vivo* studies in *Brca1* heterozygous knockout mice demonstrate that high fat diet-induced obesity leads to glucose intolerance in association with elevation in epithelial cell DNA damage and greater mammary tumor penetrance relative to mice fed a low fat diet. The data presented provide mechanistic evidence supporting an increased risk of breast cancer development in *BRCA* mutation carriers with elevated bodyweight and metabolic dysfunction, and importantly, provides clinically relevant strategies for risk reduction.

## RESULTS

### Obesity positively correlates with breast epithelial cell DNA damage in women carrying a mutation in *BRCA1* or *BRCA2*

To assess levels of DNA damage in normal breast epithelium in association with bodyweight in women carrying a *BRCA1* or *BRCA2* mutation, tissue microarrays were constructed from non-cancerous breast tissue obtained from 72 women undergoing mastectomy. The study population included *BRCA1* (n=43) and *BRCA2* (n=29) mutation carriers who had documented body mass index (BMI, kg/m^2^) between 17.7 and 44.9 (median 23.7) at the time of surgery as shown in **Table 1**. When grouping the population by BMI category of lean (BMI ≤ 24.9 kg/m^2^, n=46) or overweight/obese (BMI ≥ 25.0 kg/m^2^, n=26), median age is significantly higher in the overweight/obese group compared to the lean group (44.5 vs 38.5, respectively, *P*=0.01). Additional clinical features elevated in the overweight/obese group compared to the lean group include percent of subjects diagnosed with dyslipidemia (23.1% vs 2.2%, *P*=0.01) and with hypertension (23.1% vs 4.3%, *P*=0.02). The lean group also has a greater representation of pre-menopausal vs post-menopausal subjects compared to the overweight/obese group (*P*=0.04). Diagnosis of diabetes, race, presence of invasive tumor, tumor subtype and *BRCA*1 vs *BRCA*2 mutation were not significantly different between the two BMI groups (**Table 1**).

**Table 1.** Baseline characteristics of study population based on BMI category.

| Variables | All (n = 72) | Lean (n = 46) | Overweight/Obese (n = 26) | P |
| --- | --- | --- | --- | --- |
| BMI, median (range) | 23.7 (17.7-44.9) | 21.8 (17.7-24.7) | 28.8 (25.3-44.9) | <0.01 |
| BRCA mutation, n (%) |  |  |  | 0.46 |
| BRCA1 | 42 (58.3%) | 29 (63.0%) | 14 (53.8%) |  |
| BRCA2 | 30 (41.7%) | 17 (37.0%) | 12 (46.2%) |  |
| Age, median (range) | 40 (25-69) | 38.5 (25-60) | 44.5 (28-69) | 0.01 |
| Diabetes, n (%) |  |  |  | 0.36 |
| No | 71 (98.6%) | 46 (100.0%) | 25 (96.2%) |  |
| Yes | 1 (1.4%) | 0 (0%) | 1 (3.8%) |  |
| Dyslipidemia, n (%) |  |  |  | 0.01 |
| No | 65 (90.3%) | 45 (97.8%) | 20 (76.9%) |  |
| Yes | 7 (9.7%) | 1 (2.2%) | 6 (23.1%) |  |
| Hypertension, n (%) |  |  |  | 0.02 |
| No | 64 (88.9%) | 44 (95.7%) | 20 (76.9%) |  |
| Yes | 8 (11.1%) | 2 (4.3%) | 6 (23.1%) |  |
| Menopausal status, n (%) |  |  |  | 0.037 |
| Pre- | 48 (66.7%) | 35 (76.1%) | 13 (50.0%) |  |
| Post- | 24 (33.3%) | 11 (23.9%) | 13 (50.0%) |  |
| Race, n (%) |  |  |  | 0.19 |
| Asian | 1 (1.4%) | 1 (2.2%) | 0 (0.0%) |  |
| Black | 2 (2.8%) | 2 (4.3%) | 0 (0.0%) |  |
| Other | 2 (2.8%) | 0 (0.0%) | 2 (7.7%) |  |
| White | 59 (81.9%) | 36 (78.3%) | 23 (88.5%) |  |
| Missing | 8 (11.1%) | 7 (15.2%) | 1 (3.8%) |  |
| Invasive tumor present, n (%) |  |  |  | 1 |
| No | 40 (55.6%) | 26 (56.5%) | 14 (53.8%) |  |
| Yes | 32 (44.4%) | 20 (43.5%) | 12 (46.2%) |  |
| Tumor subtype, n(%) |  |  |  | 1 |
| HR+ | 23 (31.9%) | 15 (32.6%) | 8 (30.8%) |  |
| HER2+ | 1 (1.4%) | 1 (2.2%) | 0 (0.0%) |  |
| TNBC | 10 (13.9%) | 6 (13.0%) | 4 (15.4%) |  |
| N/A | 38 (52.8%) | 24 (52.2%) | 14 (53.8%) |  |

Abbreviations: BMI, body mass index; HR, hormone receptor; HER2, human epidermal growth factor receptor 2; TNBC, triple negative breast cancer

Immunofluorescence staining for the DNA double strand break marker γH2AX was performed with nuclear counterstain Hoechst to visualize the number of foci of DNA damage per epithelial cell (**Fig. 1A**). Among *BRCA1* and *BRCA2* mutation carriers, BMI was positively associated with breast epithelial cell DNA damage as quantified by # of γH2AX foci/100 cells (**Fig. 1B**). Age was also found to be significantly correlated with DNA damage (**Fig. 1C**). While this correlation diminished when adjusting for BMI (*P*=0.11, **Table 2**), BMI remained positively associated with DNA damage when adjusting for age (*P*=0.025, **Table 2**). Post-menopausal women were found to exhibit significantly higher levels of DNA damage compared to pre-menopausal women (**Fig. 1D**). Additionally, circulating levels of sex hormone binding globulin (SHBG), which binds estrogen to decrease its bioavailability, were negatively correlated with breast epithelial cell DNA damage (**Fig. 1E**). This negative association remains significant when adjusting for both BMI and age (*P*=0.047 and *P*=0.026, respectively, **Table 2**). Elevated BMI is often coupled to insulin resistance, a hallmark of metabolic dysfunction. Accordingly, fasting serum levels of insulin and HOMA2 IR were positively correlated with levels of breast epithelial cell DNA damage while glucose was not (**Fig. 1F-H**). Insulin and HOMA2 IR retained significance after adjustments for either BMI or age (*P*<0.001 for both, **Table 2**). No correlation with DNA damage was observed for circulating biomarkers of inflammation including high-sensitivity C-reactive protein (hsCRP) and interleukin-6 (IL-6) or with crown-like structures (CLS), a histological marker of local breast adipose inflammation (*30*) (**Fig 1. I-K**). These data indicate that among *BRCA* mutation carriers, elevated bodyweight is a risk factor for breast epithelial cell DNA damage. Furthermore, specific obesity-associated factors including insulin resistance and estrogen balance may be important drivers of this risk.

**Fig. 1.**
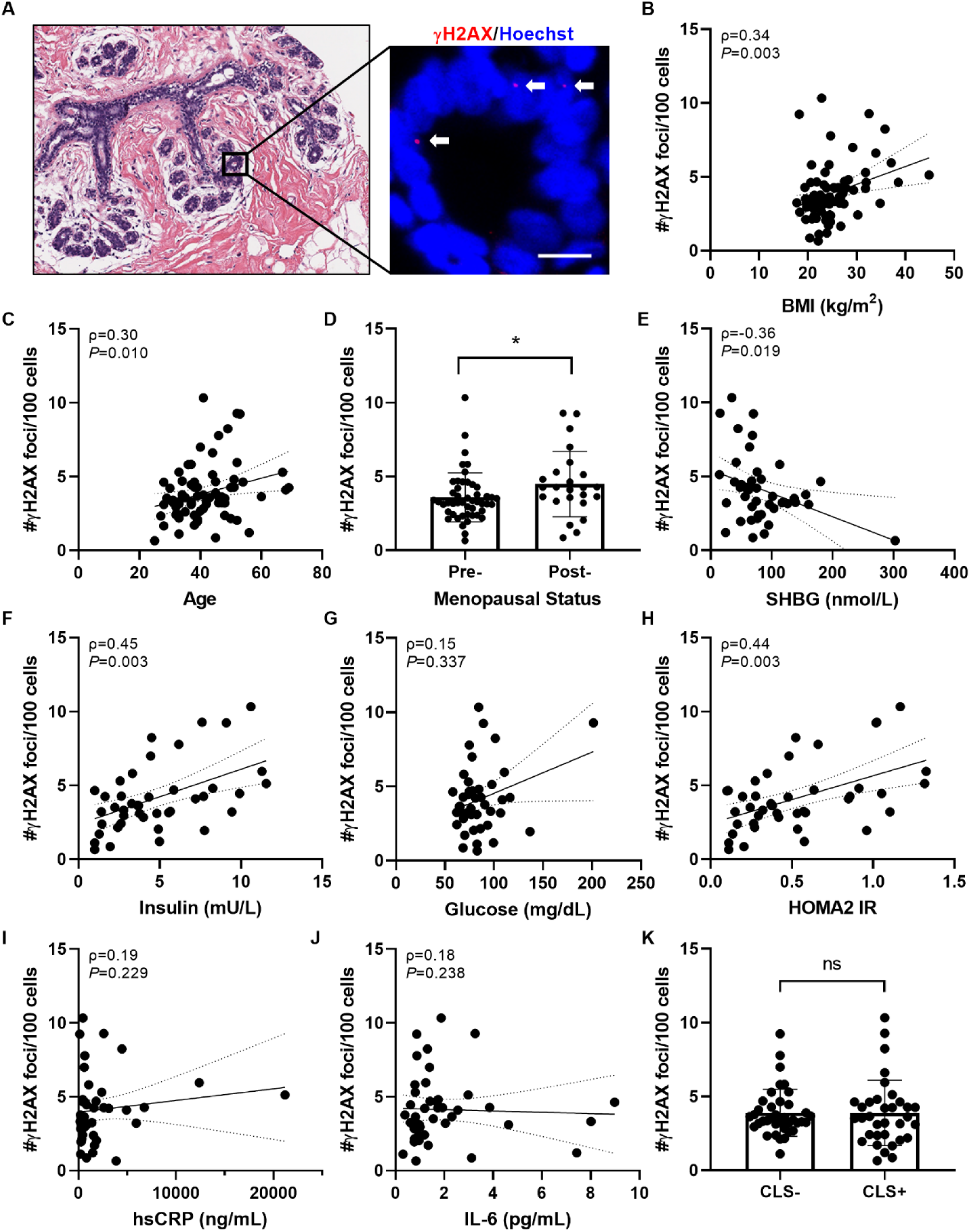
BMI and additional clinical characteristics are positively correlated with DNA damage in breast epithelium of women carrying a *BRCA* mutation. (**A**) Representative image of tissue microarray section of normal breast epithelium shown by H&E stain (left) and by IF staining (right) for γH2AX (red, arrows) co-localizing with Hoechst (blue), scale bar=10µM. (**B-C**) Correlation between epithelial cell DNA damage as measured by #γH2AX foci/100 cells with clinical characteristics including BMI and age. (**D**) Average DNA damage in the study population grouped by menopausal status: pre-menopausal, n=48 and post-menopausal, n=24. (**E-J**) Epithelial cell DNA damage correlated with circulating serum biomarkers in a subset of the study population with available fasting serum at the time of surgery (n=43). (**K**) Average DNA damage in the study population when grouped by those exhibiting histological breast adipose tissue inflammation defined as presence of crown-like structures (CLS) vs those with no CLS present (i.e. CLS- vs CLS+). Two-tailed Mann Whitney test was used to determine significant differences in grouped comparisons and data is presented as mean +/- SD. and Correlation between variables were assessed by Spearman’s rank correlation coefficient (ρ). Associated *P* value and ρ are shown for continuous variables with 95% confidence intervals. \**P*<0.05; ns, not significant; n=72 unless otherwise stated.

**Table 2.** Association of clinical features and blood biomarkers with DNA damage, adjusting for age or BMI.

| Variables | Correction | P | Correction | P |
| --- | --- | --- | --- | --- |
| BMI |  |  | Age | 0.025 |
| Age | BMI | 0.115 |  |  |
| SHBG (nmol/L) | BMI | 0.047 | Age | 0.026 |
| Insulin (mU/L) | BMI | <0.001 | Age | <0.001 |
| HOMA2 IR | BMI | <0.001 | Age | <0.001 |

Abbreviations: BMI, body mass index; SHBG, steroid hormone binding globulin, HOMA2 IR, homeostatic model assessment 2 for insulin resistance

### Elevated bodyweight is associated with significant differences in gene expression in breast adipose tissue and in breast epithelial cells of *BRCA* mutation carriers

To identify changes associated with obesity in breast epithelial cells and in the breast adipose microenvironment that may be linked to DNA damage, we conducted RNA-seq studies on both isolated primary breast epithelial cells and non-cancerous whole breast tissue obtained from *BRCA1* and *BRCA2* mutation carriers.

RNA-seq was conducted on breast tissue pieces obtained from lean (BMI ≤ 24.9kg/m^2^, n=64) and overweight/obese (BMI ≥25 kg/m^2^, n=67) *BRCA* mutation carriers. An unsupervised heatmap was constructed which shows general clustering of lean cases and clustering of overweight/obese cases by gene expression (**Fig. 2A**). 2329 genes were significantly upregulated by obesity and 1866 were significantly downregulated. Ingenuity Pathway Analysis (IPA) identified several pathways that were significantly altered in the overweight/obese cases which include pathways associated with obesity and metabolic dysfunction, such as “Phagosome Formation”, “LXR/RXR Activation”, “Tumor Microenvironment Pathway Activation”, and “Estrogen Biosynthesis” (**Fig. 2B**). A heatmap of genes involved in estrogen regulation shows a significant increase in many genes involved in the bioactivity, biosynthesis and activation of estrogens, including steroid sulfatase, 3βHSD1, AKR1C3, AKR1B15, 17βHSD1 and aromatase (CYP19A1) (**Fig. 2C**). Conversely, gene expression of 17βHSD8, involved in estrogen inactivation, was significantly lower in overweight/obese relative to lean cases. Moreover, there were mixed effects of obesity on the expression of genes involved in estrogen catabolism to hydroxylated metabolites and neutralization by COMT.

**Fig. 2.**
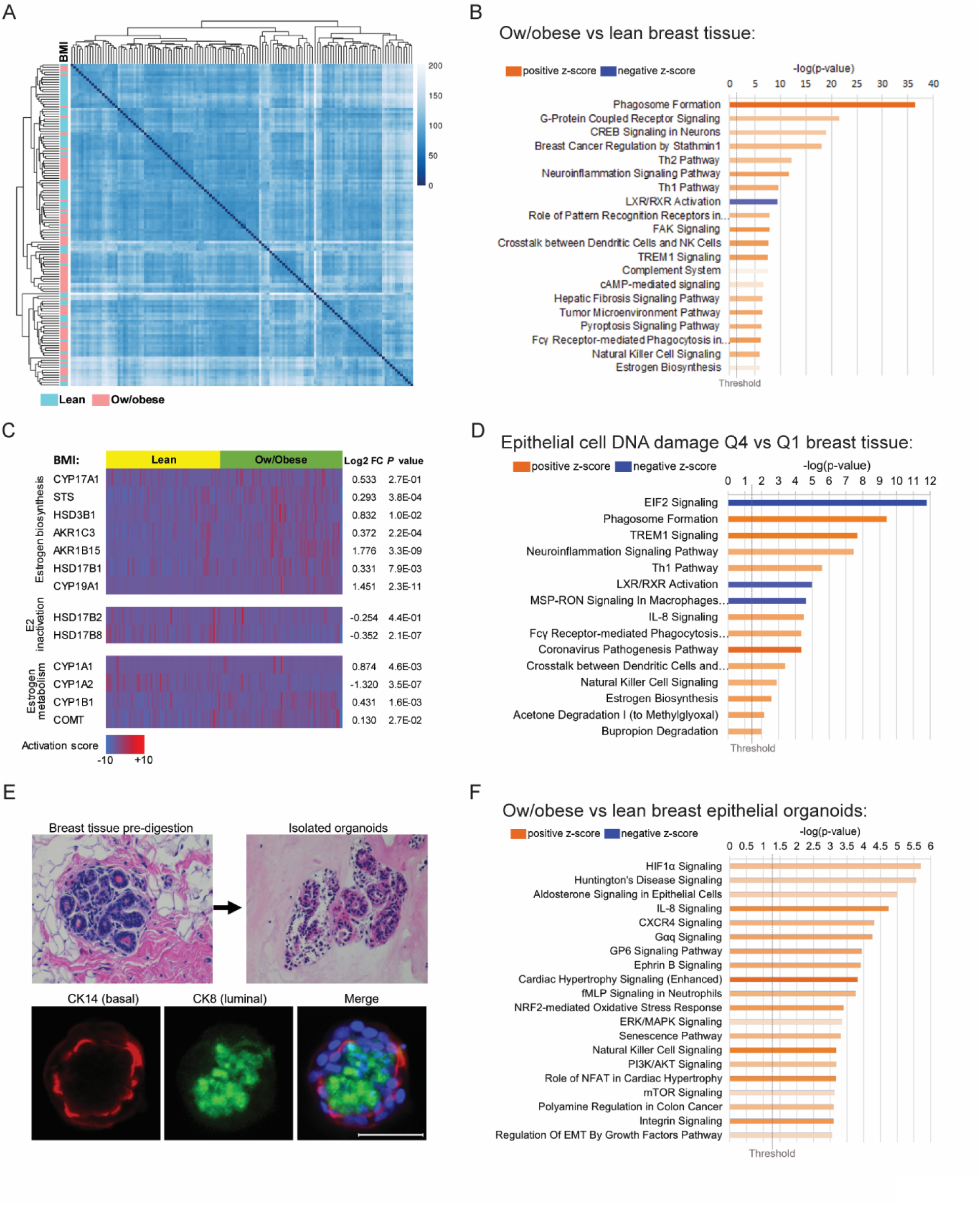
Elevated bodyweight is associated with significant changes in gene expression in breast adipose tissue and in breast epithelial cells of *BRCA* mutation carriers. (**A**) Unsupervised heatmap of whole breast tissue gene expression by RNA-seq in *BRCA* mutation carriers identified by BMI category of lean (n=64, blue) or overweight/obese (n=67, pink). (**B**) IPA analysis of RNA-Seq data showing activation (z-score) of the top 20 canonical pathways regulated in breast tissue from overweight/obese *BRCA* mutation carriers compared to lean carriers with an absolute value z-score of >0.5. (**C**) Heatmap of RNA-seq gene expression data generated from breast tissue of *BRCA* mutation carriers grouped by BMI category of lean (yellow) or overweight/obese (Ow/obese, green) showing selected genes associated with estrogen biosynthesis, estradiol (E2) inactivation, and estrogen metabolism. Corresponding gene expression (log2FC) and *P* values are shown in Ow/obese relative to lean tissue. (**D**) DNA damage in breast epithelial cells was quantified in tissue sections from n=61 patients from whom corresponding whole breast tissue RNA-seq data was also available. The cases were stratified by quartile of DNA damage and the breast tissue gene expression from cases with the highest level of DNA damage (quartile 4, Q4) were compared to cases with the lowest level (quartile 1, Q1) of DNA damage. Top 15 canonical pathways regulated in Q4 vs Q1 with an absolute value z-score of >2.0 are shown. (**E**) Representative H&E-stained images of a breast tissue section before digestion and epithelial organoids after isolation are shown. Organoids stain positively for luminal marker cytokeratin 8 (CK8, green) and basal marker cytokeratin 14 (CK14, red) as shown by IF staining merged with Hoechst (blue). Scale bar= 50µM. (**F**) IPA analysis of RNA-seq gene expression data showing activation of the top 20 canonical pathways regulated in primary breast epithelial organoids from of overweight/obese (Ow/obese) *BRCA* mutation carriers (n=9) relative to lean carriers (n=10) with an absolute value z-score of >1.0 is shown. The length of the bars on all canonical pathway graphs are determined by the Fisher’s Exact Test *P* value with entities that have a -log(p-value) >1.3 shown.

To explore which changes in the breast microenvironment are associated with DNA damage in breast epithelial cells, we analyzed breast tissue pathway changes in relation to level of epithelial cell DNA damage quantified by γH2AX immunofluorescence staining (**Fig. 2D**; n=61). The level of epithelial cell DNA damage in each case was stratified by quartiles and breast tissue gene expression was compared in the highest quartile (Q4) relative to the lowest quartile (Q1), independent of BMI . The top 15 canonical pathways activated in Q4 vs Q1 breast tissue are shown (**Fig. 2D**) with several pathways being common to both DNA damage and BMI analyses (**Fig. 2D** vs **Fig. 2B**). Although the estrogen biosynthesis pathway was found to be activated in tissue from overweight/obese compared to lean cases (**Fig. 2B**, z-score=0.775, *P* value=1.14×10^-6^), a stronger activation score is found when comparing Q4 vs Q1 (**Fig. 2D**, z-score=2.646, *P* value=2.7×10^-3^), suggesting that tissue estrogen biosynthesis is highly correlated with level of breast epithelial cell DNA damage, irrespective of BMI.

Breast epithelial organoids were isolated from *BRCA* mutation carriers who were either lean (n=10) or overweight/obese (n=9) at the time of surgery. To validate and characterize the isolated epithelial organoids, immunofluorescence staining was conducted for cytokeratin 8 (CK8) and cytokeratin 14 (CK14), characteristic markers of luminal and basal epithelial cells, respectively, that are known to comprise the breast epithelium (**Fig. 2E**). 1144 genes were significantly upregulated in the setting of overweight/obesity and 537 were significantly downregulated compared to lean organoids. The top 20 canonical pathways identified by IPA as regulated in the overweight/obese organoids are shown (**Fig. 2F**) and include activation of pathways known to be associated with obesity, including “HIF1α signaling”, “IL-8 signaling”, “ERK/MAPK signaling”, and “PI3K/AKT signaling”, among others.

Collectively, these RNA-seq studies show that *BRCA1* and *BRCA2* mutation carriers who are overweight or obese have significantly altered breast epithelial cell and breast adipose microenvironment gene expression compared with lean counterparts. Moreover, our data provide rationale for further exploring whether estrogen is a driver of DNA damage in the breast.

### Crosstalk between epithelial cells and the breast adipose microenvironment

Given the significant gene expression changes identified in *BRCA* heterozygous breast adipose tissue and in breast epithelial cells in association with overweight/obesity, we next investigated whether the breast adipose microenvironment drives gene expression in breast epithelial cells. IPA Upstream Regulator tool was used to identify regulators of gene expression differences in overweight/obese relative to lean organoids. To highlight endogenous factors that may be responsible for driving gene expression changes, results were filtered to show the top 20 secreted factors. Among these factors, beta-estradiol (an estrogen) is the top predicted upstream regulator (**Table 3**). A number of additional predicated upstream organoid regulators are significantly upregulated in overweight/obese breast tissue, including several interleukins (IL2, IL15, and IL5), TGFβ1, CSF1, ANGPT2, and WNT5A. Some factors, such as insulin, are known to be elevated in obesity, but not produced locally in breast tissue and therefore do not have an observed tissue gene expression level. These data suggest that some endogenously produced factors in the overweight/obese breast microenvironment may interact with neighboring breast epithelial cells to induce gene expression changes and DNA damage.

**Table 3.** Predicted upstream regulators of gene expression differences in breast epithelial organoids from overweight/obese *BRCA* mutation carriers relative to lean carriers and associated gene expression in whole breast tissue.

| Organoid upstream regulator | Predicted activation state | Activation z-score | P-value of overlap | Breast tissue log2FC | P-value |
| --- | --- | --- | --- | --- | --- |
| beta-estradiol | Activated | 4.728 | 2.2E-10 | see Fig. 2C |  |
| IL2 | Activated | 3.402 | 3.1E-02 | 0.563 | 2.3E-01 |
| GDF2 | Activated | 3.217 | 4.9E-03 | -0.081 | 9.8E-01 |
| IL15 | Activated | 3.152 | 1.5E-03 | 0.299 | 4.1E-05 |
| TNFSF11 | Activated | 3.125 | 3.2E-02 | -0.757 | 9.7E-02 |
| Insulin | Activated | 3.113 | 6.1E-03 |  |  |
| IL4 | Activated | 3.016 | 1.9E-03 | -0.25 | 7.4E-01 |
| TGFB1 | Activated | 2.942 | 6.0E-09 | 0.455 | 2.2E-08 |
| hydrogen peroxide | Activated | 2.839 | 3.1E-03 |  |  |
| IL3 | Activated | 2.674 | 7.6E-04 | -0.122 | 9.7E-01 |
| CSF1 | Activated | 2.602 | 8.9E-03 | 0.35 | 1.4E-06 |
| Lh | Activated | 2.598 | 1.7E-03 |  |  |
| dinoprost (PGF2α) | Activated | 2.569 | 2.9E-02 | | |
| IL5 | Activated | 2.496 | 5.5E-03 | 0.173 | 6.9E-01 |
| ATP | Activated | 2.443 | 8.7E-03 |  |  |
| MDK | Activated | 2.433 | 2.9E-02 | -0.34 | 4.4E-03 |
| AGT | Activated | 2.345 | 4.2E-03 | -0.56 | 9.6E-04 |
| ANGPT2 | Activated | 2.329 | 1.1E-03 | 0.38 | 9.2E-05 |
| WNT5A | Activated | 2.292 | 1.7E-03 | 0.184 | 1.3E-01 |
| pyruvic acid | Activated | 2.156 | 1.5E-03 |  |  |

### Targeting estrogen in breast tissue from *BRCA* mutation carriers reduces epithelial cell DNA damage

Next, we conducted mechanistic studies to determine whether targeting estrogen signaling or biosynthesis in breast tissue would lead to decreased levels of breast epithelial cell DNA damage. We first conducted immunohistochemistry (IHC) staining to verify that normal epithelium from *BRCA1* and *BRCA2* mutation carriers express the estrogen receptor (ERα). Epithelial cells staining positively for ERα were found throughout the epithelium among carriers of *BRCA1* or *BRCA2* mutations (representative images shown in **Fig. 3A, top row**). IF staining was then conducted to visualize whether γH2AX foci co-localize with ERα positive cells. Representative images are shown which highlight ERα positive cells frequently staining positively for γH2AX foci (**Fig. 3A, bottom row**). Next, we tested whether disrupting estrogen signaling through use of the drug fulvestrant, which degrades the estrogen receptor, would impact levels of DNA damage in the breast. Breast tissue was obtained from *BRCA* mutation carriers undergoing surgery (n=7) and were plated as explants in the presence of fulvestrant (100µM) or vehicle for 24 hours (**Fig. 3B**). Explants were formalin fixed and sectioned for assessment of breast epithelial cell DNA damage by IF staining. An approximately 32.5% reduction in DNA damage was observed overall after treatment with fulvestrant (**Fig. 3C**).

**Fig. 3.**
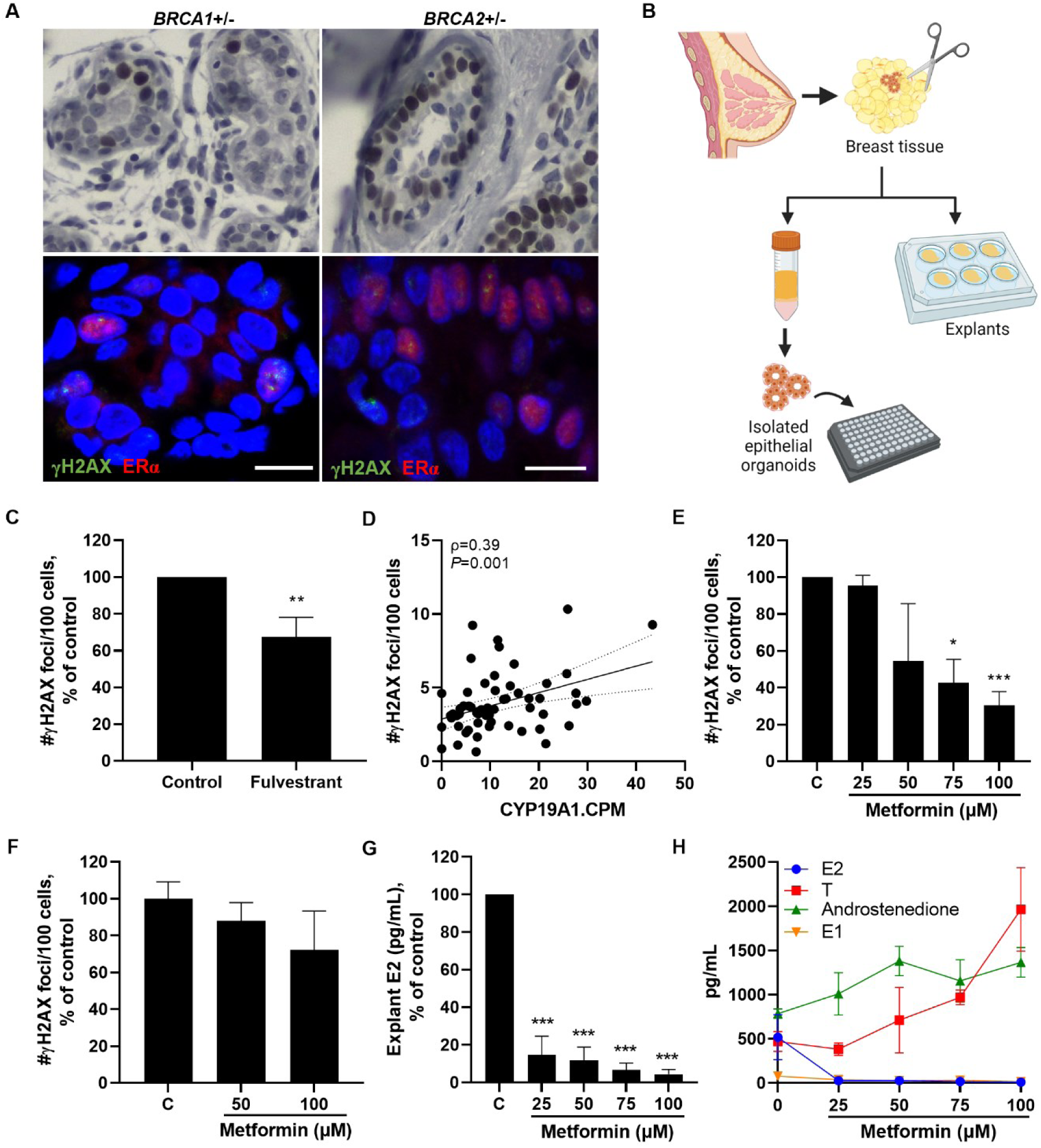
Targeting estrogen signaling or production in breast tissue decreases epithelial cell DNA damage in in women carrying a mutation in *BRCA1* or *BRCA2*. (**A**) Representative IHC staining of ERα expression in breast epithelium from carriers of a *BRCA1* or *BRCA2* mutation (top panel). Representative IF staining showing co-localization of #γH2AX foci (green) with ERα positive cells (red) (bottom panel), scale bar=10µM. (**B**) Experimental schematic showing collection of breast tissue and plating of explants or isolation of primary breast epithelial organoids for treatment studies. (**C**) Breast epithelial cell DNA damage assessed by IF (#γH2AX foci/100 cells) in e*x vivo* breast adipose tissue explants from *BRCA* mutation carriers treated with fulvestrant (100nM) for 24 hours (pooled average of n=7 patients). (**D**) Aromatase expression in breast tissue from *BRCA* mutation carriers (RNA-seq counts per million, CPM) correlated with level of breast epithelial cell DNA damage in corresponding tissue sections (n=61). Spearman’s rank correlation coefficient (ρ) and associated *P* value are shown with 95% confidence intervals. (**E**) Breast epithelial cell DNA damage in e*x vivo* breast adipose tissue explants from *BRCA* mutation carriers treated with metformin (0-100µM) for 24 hours (pooled average of n=3 patients). (**F**) DNA damage in isolated primary breast epithelial cells from *BRCA* mutation carriers treated with metformin (0-100µM) for 24 hours (representative of n=2 experiments). (**G**) Average 17β-estradiol (E2) levels and (**H**) overlay of E2, testosterone (T), androstenedione, and estrone (E1) levels in e*x vivo* breast adipose explants after 24-hour treatment with metformin (pooled average of n=3 patients). Student’s t-test was used to determine significant differences from control unless otherwise stated. Data is presented as mean +/- SEM. \**P* <0.05, \*\**P* <0.01, \*\*\**P*<0.001.

Next, we hypothesized that targeting estrogen biosynthesis in the breast by downregulating aromatase expression would lead to less estrogen exposure to the epithelial cells, and consequently decreased DNA damage. In support of this hypothesis, RNA-seq data from *BRCA1* and *BRCA2* mutation carriers showed a positive correlation between breast adipose aromatase expression and level of breast epithelial cell DNA damage (**Fig. 3D**). Importantly, since aromatase expression is known to be upregulated in association with obesity, we conducted additional statistical analyses to adjust for BMI and found that aromatase remained independently positively correlated with DNA damage (*P*=0.037). To target estrogen biosynthesis, we utilized metformin, a widely used antidiabetic drug which has also been shown to decrease aromatase production in the breast via stimulation of AMP-activated protein kinase (AMPK) in adipose stromal cells (*31, 32*). Breast tissue obtained from *BRCA* mutation carriers (n=3) were plated as explants and treated with metformin (0-100µM) for 24 hours followed by IF assessment of breast epithelial cell DNA damage. A dose-dependent decrease in DNA damage was observed with significant differences after 75 and 100µM of metformin treatment (**Fig. 3E**). Since metformin is known to decrease aromatase expression in adipose stromal cells surrounding breast epithelial cells, we digested breast tissue to isolate the epithelial cells from their microenvironment (**Fig. 3B**) and treated them with metformin for 24 hours to determine if the presence of the breast microenvironment is required for the effect of metformin on DNA damage. Although there was a modest trend for reduction in DNA damage with increasing doses of metformin, these results were not significant (**Fig. 3F**). Consistently, tissue levels of estradiol (E2) were markedly reduced in breast explants after 24-hour metformin treatment in a dose-dependent manner (**Fig. 3G**). Additionally, testosterone and androstendione, which are converted to E2 and estrone (E1) by aromatase, respectively, were increased in explants following treatment with metformin while both E1 and E2 decreased (**Fig. 3H**). These data show that metformin treatment leads to decreased estrogen biosynthesis in breast tissue in association with reduction in epithelial cell DNA damage.

### Local and systemic factors contribute to DNA damage in *BRCA1* and *BRCA2* heterozygous breast epithelial cells

Our data support a paracrine interaction between adipose tissue and breast epithelial cells. Having found a direct role for estradiol in mediating DNA damage in primary human tissues, we next explored the role of additional obesity-associated factors, including those present in breast adipose tissue conditioned media (CM), as well as recombinant leptin and insulin. To first investigate whether factors derived from breast adipose tissue have the ability to directly induce DNA damage in *BRCA* mutant breast epithelial cells, *BRCA1* heterozygous knockout MCF-10A cells were treated with CM from reduction mammoplasty or non-tumor quadrants of mastectomy tissue (**Fig. 4A**, n=36, BMI: 20.6-40.1 kg/m^2^). Breast adipose CM treatment was positively correlated with DNA damage as a function of the patient’s BMI, as measured by immunofluorescence of γH2AX foci (**Fig. 4B**). RNA-seq was performed in a subset of CM-treated samples (lean and obese, n=3/group). Results demonstrate that consistent with DNA damage measurements (**Fig. S1**), IPA analysis of differentially regulated genes in the obese CM treated cells relative to lean showed increased activation of functions associated with DNA damage and genomic instability including “Formation of micronuclei”, “Chromosomal instability”, and “Breakage of chromosomes”. Alternatively, activation of functions associated with DNA repair were decreased, including “Repair of DNA” and “Checkpoint control” (**Table 4**).

**Fig. 4.**
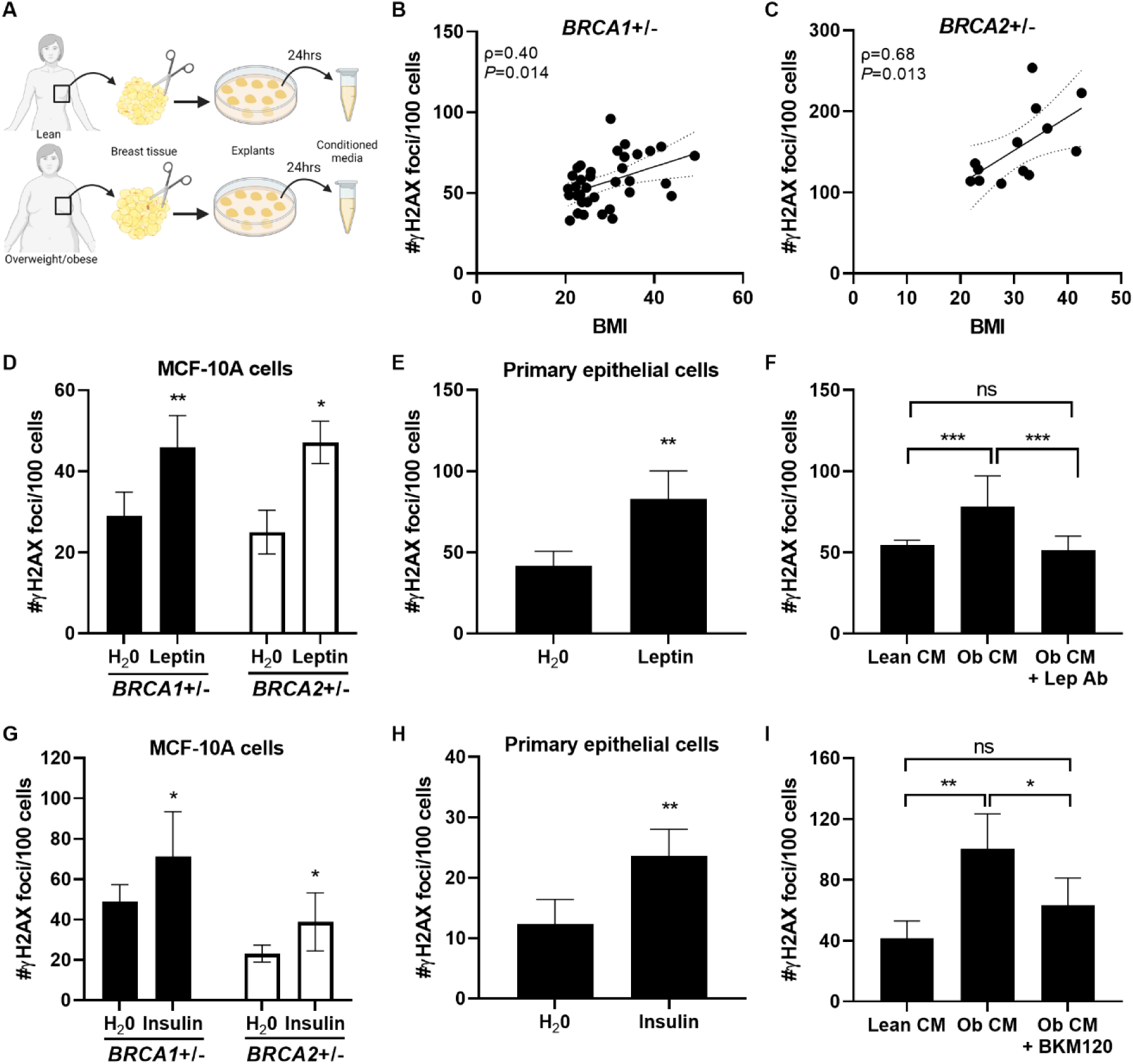
Obesity-induced changes to the local breast adipose microenvironment promotes DNA damage in *BRCA1* and *BRCA2* heterozygous breast epithelial cells. (**A**) Experimental schematic showing the collection of breast adipose tissue conditioned media (CM) from lean and overweight/obese women. (**B**) MCF-10A cells were treated with CM for 24 hours. DNA damage assessed by IF (#γH2AX foci/100 cells) is shown correlated with BMI in *BRCA1*+/- (n=36 CM cases) and (**C**) *BRCA2*+/- (n=13 CM cases) MCF-10A cells. Spearman’s rank correlation coefficient (ρ) and associated *P* value are shown along with 95% confidence intervals. (**D**) DNA damage in *BRCA1*+/- and *BRCA2*+/- MCF-10A cells and in (**E**) primary *BRCA1*+/- breast epithelial cells treated with leptin (400ng/µl) for 24 hours. (**F**) DNA damage in *BRCA1*+/- MCF-10A cells after 24-hour treatment with lean CM, obese (ob) CM, or ob CM in the presence of a leptin neutralizing antibody (Lep Ab). (**G**) DNA damage in *BRCA1*+/- and *BRCA2*+/- MCF-10A cells and in (**H**) primary *BRCA2*+/- breast epithelial cells treated with insulin (100nM) for 24 hours. (**I**) DNA damage in *BRCA1*+/- MCF-10A cells after 24-hour treatment with lean CM, ob CM, or ob CM in the presence of PI3K inhibitor BKM120 (1µM). Student’s t-test was used to determine significant differences in (**D-I)**. All experiments in MCF-10A cells were conducted a minimum of two times with representative results from one experiment shown. Data in primary cells were generated from cells treated in triplicate. Data is presented as mean +/- SD. \**P* <0.05, \*\**P* <0.01, \*\*\**P* <0.001, ns= not significant.

**Table 4.** Activation of diseases or functions associated with DNA damage or DNA repair in *BRCA1*+/- epithelial cells treated with breast adipose tissue condition media derived from obese women relative to lean women.

| Categories | Diseases or functions annotation | P-value | Predicted activation state | Activation z-score | # Molecules |
| --- | --- | --- | --- | --- | --- |
| Cellular Assembly and Organization | Formation of micronuclei | 2.53E-06 | Increased | 2.756 | 9 |
| DNA Replication, Recombination & Repair | Chromosomal aberration | 5.37E-06 | Increased | 2.853 | 31 |
|  | Chromosomal instability | 2.43E-08 | Increased | 2.603 | 19 |
|  | Breakage of chromosomes | 2.88E-05 | Increased | 2.488 | 11 |
| Cell Cycle; DNA Replication, Recombination & Repair | Checkpoint control | 1.99E-06 | Decreased | -2.756 | 15 |
|  | Spindle checkpoint | 9.33E-07 | Decreased | -2.035 | 12 |
| DNA Replication, Recombination & Repair | Repair of DNA | 4.36E-09 | Decreased | -3.334 | 47 |
|  | Double-stranded DNA break repair of tumor cell lines | 9.94E-06 | Decreased | -2.241 | 14 |
|  | Metabolism of DNA | 2.10E-10 | Decreased | -2.09 | 54 |

To determine whether effects of CM on DNA damage were generalizable to *BRCA2* mutation carriers, a subset of CM cases were tested in MCF-10A cells carrying a heterozygous *BRCA2* mutation, generated using CRISPR-Cas9 gene editing (see Supplementary Materials and Methods). A positive correlation between BMI and DNA damage was also observed in these cells (**Fig. 4C**). These studies demonstrate that factors secreted by breast adipose tissue directly stimulate DNA damage in breast epithelial cells. Furthermore, given the lack of estrogen receptor expression in MCF-10A cells (*33*), these studies also highlight the existence of additional factors beyond estrogen that may be contributing to DNA damage induction in the setting of obesity in *BRCA1* and *BRCA2* mutant breast epithelial cells.

The expression of leptin, known to be directly correlated with adiposity, is significantly higher in overweight/obese compared to lean breast tissue from *BRCA* mutation carriers (Table S1, log2FC= 0.61, *P*=3.48×10^-6^). A number of studies have found leptin to have pro-mitogenic and anti-apoptotic effects in breast cancer cells (*34–37*). However, its effects on normal breast epithelial cells are less well characterized. Here, we treated both *BRCA1* and *BRCA2* heterozygous MCF-10A cells with leptin (400ng/mL) for 24 hours and found a significant induction of DNA damage in both cell lines (**Fig. 4D**) and in primary breast epithelial cells (**Fig. 4E**). Additionally, the ability of obese CM to induce DNA damage in *BRCA1* heterozygous breast epithelial cells is blocked when treating in the presence of a leptin neutralizing antibody (**Fig. 4F**).

Next, having identified insulin as positively correlated with DNA damage in tissue microarrays from *BRCA* mutation carriers, independent of BMI (**Fig. 1F**, **Table 2**), and as a top upstream regulator of gene expression in primary breast epithelial organoids from overweight/obese women (**Fig. 3A**), we conducted additional mechanistic studies to determine whether insulin can directly induce DNA damage. Treatment of *BRCA1* and *BRCA2* heterozygous knockout MCF-10A cells with insulin (100nM) for 24 hours resulted in a significant increase in DNA damage in both cell lines (**Fig. 4G**) and in primary breast epithelial cells (**Fig. 4H**). Both leptin and insulin have been shown to act via PI3K (*38, 39*). Treatment of *BRCA1* heterozygous breast epithelial cells with a PI3K inhibitor, BKM120 (1µM), was effective at reducing obese CM-induced DNA damage (**Fig. 4I**). These data show that factors produced locally by obese breast adipose tissue or elevated with metabolic dysfunction contribute to induction of DNA damage in *BRCA* heterozygous knockout breast epithelial cells.

### High fat diet feeding is associated with elevated mammary gland DNA damage and early tumor penetrance in female *Brca1* heterozygous knockout mice

DNA damage is a known driver of chromosomal defects that can lead to cancer. However, whether obesity-associated elevation in breast epithelial cell DNA damage is linked to breast cancer penetrance in the setting of a heterozygous *BRCA* mutation has not been established. To investigate this question, we conducted preclinical studies utilizing mice that were developed to carry a whole-body heterozygous loss in *Brca1* (*Brca1*+/-) on a C57Bl/6 background. Four-week-old female *Brca1*+/- mice were randomized to receive low fat diet (LFD) or high fat diet (HFD) for 22 weeks to produce lean and obese mice, respectively (**Fig. 5A**). Mice fed HFD gained significantly more weight than LFD fed mice and weighed on average 34.1g vs 23.3g, respectively, at the time of sacrifice (**Fig. 5B**). Overall adiposity was also increased in association with HFD feeding as determined by greater accumulation of subcutaneous and visceral fat compared to the LFD group (**Fig. S2)**. To confirm that the HFD-fed mice exhibit altered metabolic homeostasis in our *Brca1*+/- model of diet-induced obesity, glucose tolerance tests were conducted after 21 weeks on experimental diets, highlighting delayed clearance of glucose from blood over 90 minutes post-intraperitoneal injection of glucose in the HFD group compared to LFD-fed mice (**Fig. 5C** **&** **D**). To determine whether changes observed in the mammary fat pad of *Brca1*+/- mice in response to feeding were analogous to those seen in the breast tissue of women in relation to obesity, RNA-seq was conducted on inguinal mammary fat pads from LFD and HFD mice harvested at sacrifice (**Table S5**). IPA was used to identify activation of the top differentially regulated canonical pathways in HFD mammary fat pads relative to LFD, results of which were juxtaposed with regulation of these same pathways in human breast tissue from overweight/obese vs lean *BRCA* mutation carriers. The top 20 canonical pathways regulated by obesity in the mouse mammary fat pad show very similar regulation patterns compared to overweight/obese human breast tissue (**Fig. 5E**), suggesting that diet-induced obesity in our *Brca1*+/- mice can serve as a model system for obesity in women carrying a *BRCA* mutation with respect to studies of the breast.

**Fig. 5.**
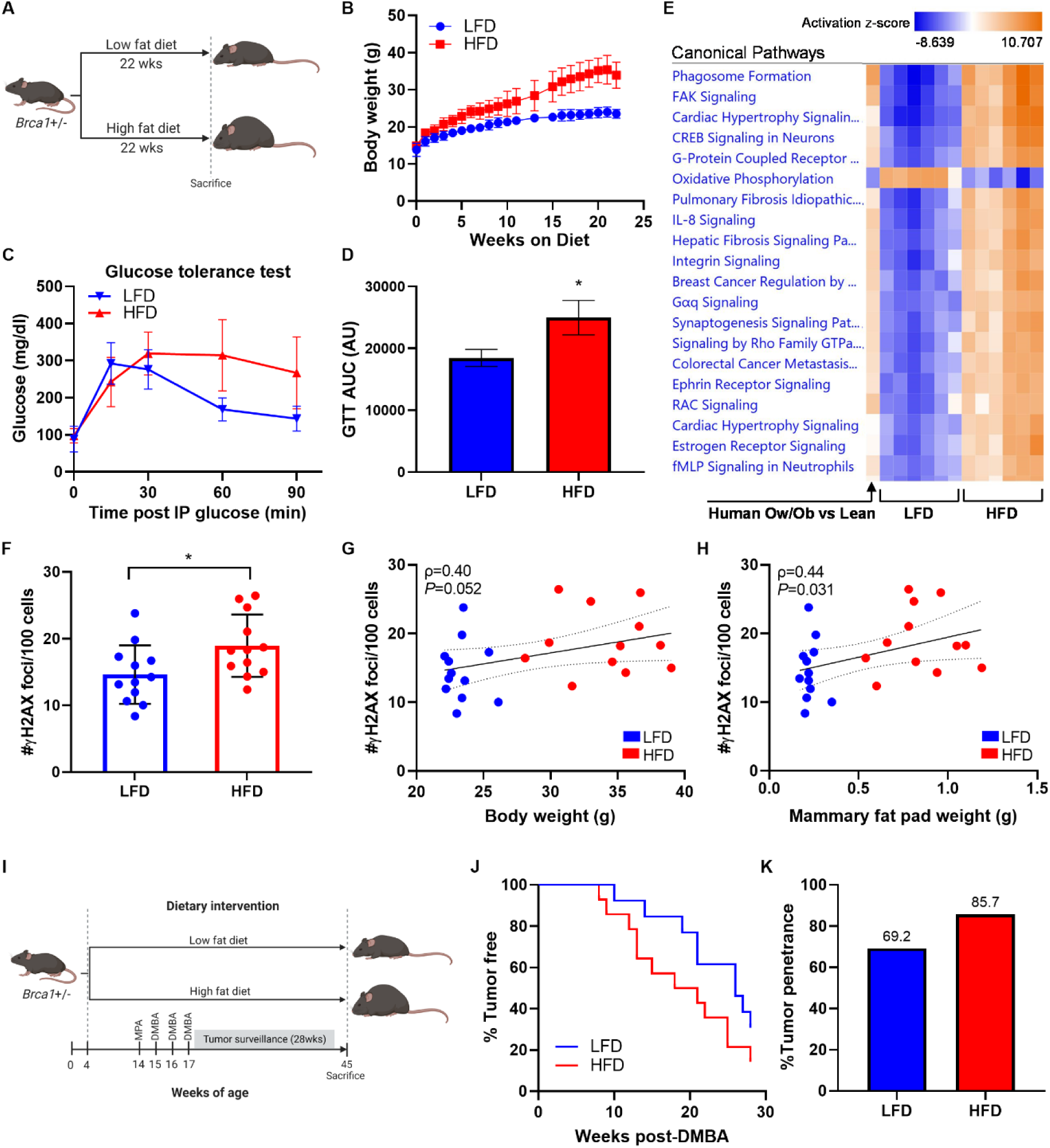
High fat diet feeding leads to elevated mammary gland DNA damage in association with increased mammary tumor penetrance and decreased tumor latency in *Brca1*+/- mice. (**A**) Experimental schematic of diet-induced obesity in female *Brca1*+/- mice (n=12/gp). (**B**) Average body weight of mice fed low fat diet (LFD) or high fat diet (HFD) over 22 wks. (**C**) Glucose tolerance test conducted one week prior to sacrifice and (**D**) area under curve (AUC) calculation for each group (mean +/- SEM). (**E**) RNA-Seq was conducted on whole mammary fat pad tissue from HFD and LFD mice (n=6/gp). Activation of top 20 canonical pathways regulated in mammary fat pads from HFD mice compared to LFD mice are shown adjacent to corresponding pathway regulation in breast tissue from overweight (Ow)/obese vs lean women carrying a *BRCA* mutation (n=64-67/gp). (**F**) DNA damage assessed by IF (#γH2AX foci/100 cells) in mammary glands at the time of sacrifice. (**G**) Correlation between mammary gland DNA damage and mouse body weight and (**H**) mammary fat pad weight among all mice. Spearman’s rank correlation coefficient (ρ) and associated *P* values are shown along with 95% confidence intervals. (**I**) Experimental schematic of MPA/DMBA-induced tumorigenesis model in female *Brca1*+/- mice randomized to LFD or HFD groups (n=13-14/gp). (**J**) Mammary tumor development in LFD and HFD mice shown as % of mice tumor free over the 28-week surveillance period. (**K**) Overall mammary tumor penetrance at the end of the surveillance period shown as % of mice in each group that developed a mammary tumor. Student’s t-test was used to determine significance unless otherwise stated. Data is presented as mean +/- SD unless otherwise stated. \**P* <0.05.

Similar to findings made in human breast tissue from *BRCA* mutation carriers, IF staining for γH2AX of *Brca1*+/- mouse mammary glands at the time of sacrifice show that HFD-fed mice have elevated levels of mammary gland DNA damage compared to LFD-fed mice (**Fig. 5F**). Furthermore, there is a trend for a positive correlation between DNA damage and bodyweight (irrespective of diet) (**Fig. 5G**) and a significant positive correlation between DNA damage and mammary fat pad weight (**Fig. 5H**), suggesting that level of adiposity may be a stronger predictor of DNA damage in mammary epithelium compared to whole body weight.

Next, we examined whether elevation in mammary gland DNA damage is associated with tumorigenesis. Female *Brca1*+/- mice were first made obese by HFD-feeding for 10 weeks and then were implanted with a subcutaneous medroxyprogesterone acetate (MPA) pellet to sensitize them to mammary tumor development upon exposure to three doses of the carcinogen 7,12-dimethylbenz[a]anthracene (DMBA) (**Fig. 5I**). Mammary tumors developed earlier in the HFD group compared with the control LFD group (**Fig. 5J**). Additionally, 85.7% of mice in the HFD group developed mammary tumors by the end of the 28-week surveillance period compared to 69.2% of mice in the LFD group (**Fig. 5K**).

### Obesity is associated with DNA damage in the fallopian tube, but not ovary, of *BRCA* mutation carriers

In addition to elevated breast cancer risk, women carrying a *BRCA1* or *BRCA2* mutation have high lifetime risk for developing ovarian cancer (*1, 2*). Since weight gain is associated with increased risk of ovarian cancer in *BRCA* mutation carriers (*40*), we extended our studies in the breast to investigate the impact of elevated BMI on DNA damage in the ovarian epithelium as well as in epithelial cells of the fallopian tube (**Fig. 6**). IF staining for γH2AX was performed with nuclear counter stain Hoechst to quantify number of foci of DNA damage per epithelial cell in non-tumorous ovarian tissue and fallopian tube fimbria from women carrying a *BRCA1* or *BRCA2* mutation undergoing prophylactic salpingo-oophorectomy. In the ovarian epithelium, there was no increase in DNA damage in the overweight/obese cases (n=12) compared to the lean cases (n=21) (**Fig. 6A**, *P*=0.59). However, there was a significant increase in DNA damage observed in the epithelial cells of the fallopian tube from overweight/obese women (n=9) compared to lean women (n=17) (**Fig. 6B**, *P*=0.03).

**Fig. 6.**
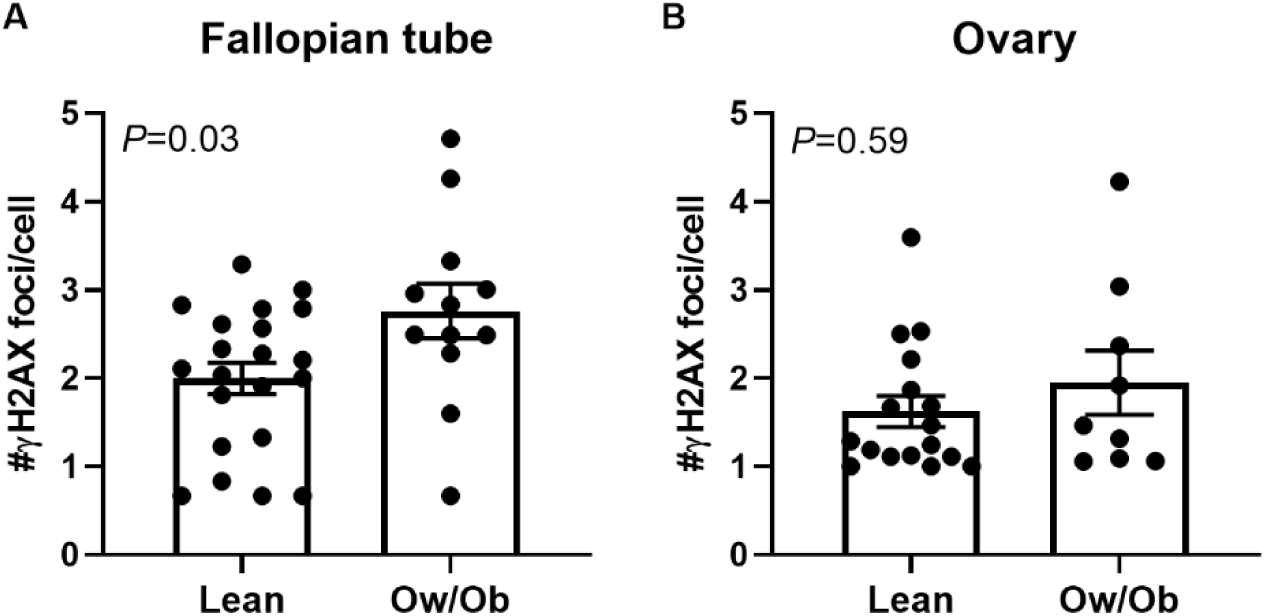
BMI is associated with DNA damage in the fallopian tube but not ovary. (**A**) DNA damage assessed by IF (#γH2AX foci/cell) in epithelial cells of the ovary and in (**B**) epithelial cells of fallopian tube fimbriae in *BRCA* mutation carriers grouped by BMI category of lean (n=17-21/gp) or overweight (Ow)/obese (Ob) (n=9-12). Two-tailed Mann Whitney test was used to determine significant differences (*P* value) between groups. Data is presented as mean +/-SEM.

## DISCUSSION

The data presented here demonstrate that BMI is positively associated with DNA damage in normal breast epithelial cells in carriers of a mutation in *BRCA1 or BRCA2*. Beyond BMI, insulin and insulin resistance, as measured by HOMA2 IR, were independently associated with DNA damage, irrespective of BMI or age. Accordingly, it is possible that *BRCA* mutation carriers who are defined as lean by BMI, but hyperinsulinemic (‘metabolically obese’), may also be at risk for elevated levels of DNA damage and consequently, breast cancer development. Although previous studies have shown that inflammation can lead to DNA damage in both normal and cancerous cells in other tissues (*41–44*), our data do not support a link between local or systemic inflammation and breast epithelial cell DNA damage.

To our knowledge, this is the first study to conduct transcriptional profiling of non-cancerous breast tissue and isolated breast epithelial cells from overweight/obese vs lean *BRCA* mutation carriers. While several factors and pathways associated with metabolic dysfunction were shown to be upregulated in breast tissue and in epithelial cells, the identification of pathways related to estrogen biosynthesis (tissue) and signaling (epithelial cells) were of particular interest given the availability of clinically approved drugs that target estrogen. Additionally, previous *in vitro* studies showed that treatment with estrogen and estrogen metabolites induced DNA damage in *BRCA1* heterozygous breast epithelial cells (*45*), providing further rationale for exploring the role of estrogen as a mediator of obesity-induced DNA damage. Here, we show that fulvestrant, an estrogen receptor degrader, is effective at reducing epithelial cell DNA damage in breast tissue explants from *BRCA* mutation carriers. However, this drug is not currently approved for use in the prevention setting and the side effects may limit its future use for this purpose. Alternatively, metformin is widely prescribed in patients with type II diabetes and has an excellent safety profile which makes this drug an intriguing option for preventative use in *BRCA* mutation carriers with excess bodyweight. We show that metformin was effective at reducing breast epithelial cell DNA damage at clinically relevant concentrations primarily due to effects on the breast adipose microenvironment. Previous studies have shown that metformin decreases adipose stromal cell expression of aromatase through activation of AMPK (*31, 32*). Our study extends these findings by demonstrating the downstream consequence of downregulation in aromatase through mass spectrometry studies which showed marked reduction in E2 in breast tissue after metformin treatment. In addition to reducing estrogen exposure, previous work has shown that metformin treatment reduces endogenous reactive oxygen species (ROS) and associated DNA damage (*46*) in a mammary epithelial cell line, providing an additional possible mechanism for the effects of metformin in our studies.

Epidemiological studies have reported decreased risk of breast cancer in *BRCA* mutation carriers in association with reduced estrogen exposure achieved via salpingo-oophorectomy surgery which diminishes ovarian estrogen production or through treatment with tamoxifen, an estrogen receptor antagonist in the breast (*47–49*). Our studies propose estrogen-mediated induction of DNA damage as a possible explanation for the protective effects observed by decreasing estrogen exposure in this population. Estrogen can induce DNA damage through various actions as reviewed by our group and others (*50, 51*), including through ligand binding to ERα which stimulates proliferation and potentially replication stress with ROS production as a byproduct of increased cellular respiration. Additionally, the metabolism of estrogen yields genotoxic metabolites, a process which produces ROS through redox cycling. These metabolites can directly interact with DNA to form adducts in an ER-independent manner. Given the multiple avenues through which estrogen can induced DNA damage in cells, additional studies are warranted to characterize the mechanisms of estrogen-induced DNA damage in breast epithelial cells from *BRCA* mutation carriers in the setting of obesity.

Interestingly, our RNA-seq analysis of *BRCA1* heterozygous MCF-10A cells treated with obese vs lean CM not only showed increased activation of pathways associated with DNA damage, but also downregulation of pathways associated with DNA repair. This raises the possibility that obesity may affect DNA repair capacity, which would be especially detrimental in cells already exhibiting defective DNA repair due to a mutation in *BRCA1* or *BRCA2*. Additional studies exploring the relationship between obesity and DNA repair would be relevant not only for *BRCA* mutation carriers, but also for the general population where obesity is associated with increased breast cancer risk in post-menopausal women (*3*). Defective DNA repair capacity would illuminate a novel mechanism through which obese non-carriers become more susceptible to breast cancer.

Our *in vitro* studies demonstrate the ability of several obesity-associated factors, including leptin and insulin, to cause DNA damage, suggesting a collective milieu of factors that may contribute to the elevation in DNA damage observed in *BRCA* mutation carriers in association with BMI. The ability of obese CM to induce damage in *BRCA1* heterozygous cells was diminished when treating in the presence of an antibody or drug that inhibits leptin or insulin signaling, respectively. Of note, since insulin signals through phosphatidylinositol 3-kinases (PI3K), we utilized BKM120, a PI3K inhibitor, to disrupt insulin actions in the presence of obese CM. It is possible that inhibiting PI3K signaling not only disrupted insulin signaling, but also signaling of other factors associated with obesity that act through PI3K, including growth factors or leptin, which collectively contributed to the observed decrease in DNA damage. Additionally, growing evidence points to a role for the PI3K pathway in the DNA damage response, however, these studies have been limited to cancer cells (*52–55*).

Our studies also show a link between obesity-induced DNA damage and tumor development using a *Brca1+/-* mouse model of diet-induced obesity. HFD-fed mice exhibited elevated mammary gland DNA damage in association with decreased latency and increased overall penetrance of mammary tumors when exposed to the carcinogen DMBA. These data suggest that the elevation in DNA damage that we observed in association with BMI in women carrying a *BRCA* mutation may also be associated with increased breast cancer penetrance. The extent to which data from this mouse model can be extrapolated to humans is somewhat limited given that we employed a carcinogen-indued tumor model, whereas in *BRCA* mutation carriers, tumors will arise after years of exposure to both endogenous and environmental factors, some of which will act as carcinogens.

Finally, our data show that obesity-associated DNA damage may not only be limited to the breast epithelium of *BRCA* mutation carriers. Although no increase in DNA damage was found in epithelial cells of the ovary in overweight/obese women undergoing prophylactic salpingo-oophorectomy, we did observe a significant increase in DNA damage in the epithelial cells of the fallopian tube in overweight/obese women. Our results are consistent with reports from recent years which point to the fallopian tube as the likely site of origin of ovarian cancer (*56, 57*), to be confirmed by ongoing clinical trials of risk-reducing salpingectomy with delayed oophorectomy, and also highlights a potential mechanism for the link between weight gain and ovarian cancer in this population.

A limitation of our study includes a cohort size of n=72 in our correlation study of DNA damage and BMI which prevented us from analyzing effects of BMI separately in *BRCA1* and *BRCA2* mutation carriers. Although both BRCA1 and BRCA2 are essential for DNA repair, their roles in the DNA damage response are not identical and each mutation is associated with different subtypes of tumor development. Larger studies assessing the relative effect of BMI on DNA damage in *BRCA1* and *BRCA2* mutation carriers separately could provide additional information to help personalize risk estimates. Additionally, levels of estrogens vary considerably during the menstrual cycle and impact proliferation of breast epithelial cells. Our studies did not account for phase of menstrual cycle when assessing DNA damage which may have led to increased variability in our data, particularly considering our identification of estrogen as a mediator of obesity-induced epithelial cell DNA damage.

Many methodological challenges exist which explain the lack of consensus in epidemiological studies attempting to ascertain modifiers of breast cancer risk in *BRCA* mutation carriers, as reviewed by Milne & Antinou (*58*). Although a number of studies have associated bodyweight with increased risk of breast cancer as discussed earlier, the largest study to date to contradict these findings showed protective effects of BMI on pre-menopausal breast cancer risk in *BRCA* mutation carriers (*11*). Drawing definitive conclusions from this study is limited due to the utilization of subject-reported BMI at the time of study questionnaire which is subject to recall bias and utilization of calculated genetic BMI score which does not necessarily predict actual observed BMI and may be influenced by dietary and environmental factors. Additionally, a subset of the overweight/obese population may have received treatment for obesity-associated co-morbidities like diabetes which potentially confounds risk assessment if these treatments or medications reduce breast cancer risk. Overall, given the inconsistencies in reported data and significant challenges in assessing modifiers of breast cancer risk in this population, the consensus to date is that there is insufficient evidence to determine the effect of bodyweight on breast cancer risk in *BRCA* mutation carriers (*58–60*). Therefore, a strength of our study is the presentation of mechanistic experimental evidence which helps to elucidate the relationship between bodyweight and breast cancer risk in this population.

Additionally, our findings provide rationale for conducting clinical trials in overweight/obese *BRCA* mutation carriers to test the efficacy of pharmacological interventions that target metabolic health, weight and/or estrogens. In fact, identifying which obesity-related factors need to be targeted for risk reduction, if not all, will have a meaningful impact on developing effective risk reduction strategies. Although recently reported results of the phase 3 randomized MA.32 trial (NCT01101438) found that addition of metformin to standard of care in non-diabetic patients with high-risk breast cancer did not significantly improve invasive disease-free survival vs placebo (*61*), it remains to be determined if metformin in the preventative setting would be effective at reducing risk of breast cancer, particularly among *BRCA* mutation carriers and those with metabolic dysfunction. Our studies point towards the potential of metformin in this setting, as it has been shown to reduce weight, as well as cause decreases in circulating levels of insulin, leptin and estrogens (*62–64*). These studies would help clarify whether accumulation of DNA damage over time is reversable or if targeted interventions prevent accumulation of further damage. Positive results would offer clinicians actionable evidence-based prevention strategies for patients in this high-risk population who opt to delay or forgo risk-reducing surgery.

## MATERIALS AND METHODS

### Study Design

The objective of this study was to gain insight into the role of obesity and metabolic dysfunction on breast cancer penetrance among carriers of germline mutations in *BRCA1* and *BRCA2* and to identify clinically relevant prevention strategies. Clinical samples including both archival tissues and prospectively collected tissues from *BRCA* mutation carriers, as well as cell lines engineered to carry a *BRCA1* or *BRCA2* heterozygous knockout mutation and *Brca1*+/- mouse models were utilized in support of this objective. All studies utilizing human tissues were conducted in accordance with protocols approved by the Institutional Review Boards of Memorial Sloan Kettering Cancer Center (MSKCC) under protocol #10-040 and Weill Cornell Medicine under protocols #1510016712, 1004010984-01 and 1612017836. Informed consent from each subject was obtained by study investigators prior to tissue collection. Animal experiments were conducted in accordance with an approved Institutional Animal Care and Use Committee protocol (#2018-0058) at Weill Cornell Medicine.

Studies utilizing archival tissues were coded and DNA damage was analyzed in a blinded fashion. Studies utilizing prospectively collected tissues and *in vitro* treatment studies were not blinded, however, DNA damage was analyzed by immunofluorescence staining using methodology to limit bias as described in the section “*Confocal microscopy & quantification of* γ*H2AX foci*” below. Sample size power calculations were performed for human breast tissue microarray construction (BMI vs DNA damage study) and in animal studies. Any sample exclusion criteria are described in the sections below or in the figure legends.

### Human breast tissue microarray construction & study population

Archival paraffin blocks of embedded non-tumorous breast tissue were obtained from 72 women carrying a *BRCA1* (n=42) or *BRCA2* (n=30) mutation who had previously undergone prophylactic or therapeutic mastectomy at MSKCC from 2011-2016. **Table 1** describes the clinical characteristics of the study population which were extracted from electronic medical records. BMI was calculated using height and weight recorded prior to surgery (kg/m^2^) and menopausal status was determined per criteria established by the National Comprehensive Cancer Network ((*65*). A pathologist reviewed hematoxylin & eosin-stained sections from each block to identify areas enriched in breast epithelium. Cores measuring 1.5mm in diameter from identified epithelial areas of each case were incorporated into paraffin blocks for the construction of tissue microarrays. Each tissue microarray was constructed with cases representing an equal distribution of clinical characteristics, including *BRCA1* or *BRCA2* mutation status and BMI. Unstained sections were cut from each tissue microarray and used for quantification of breast epithelial cell DNA damage by immunofluorescence staining as described in the section below.

### Assessment of DNA damage by immunofluorescence staining

To quantify epithelial cell DNA damage, immunofluorescence staining of the DNA double strand break marker γH2AX was conducted on human tissue sections, mouse mammary gland tissue sections, or plated cells. Antibodies/reagents that were used include: primary γH2AX (p Ser139) antibody (Novus Biologicals #NB100-74435 unless otherwise stated) at 1:300 dilution, Goat anti-Mouse Alexa Fluor 546 secondary antibody (Life technologies #A11030) at 1:1000 dilution, Hoechst 33342 nuclear stain (Santa Cruz Biotechnology #SC-495790) at 1:1000 dilution, CAS block (Life Technologies #008120), M.O.M (Mouse-on-Mouse) immunodetection kit (Vector Laboratories # BMK-2202), and ProLong Gold Antifade Mountant (Invitrogen # P36934). Full staining procedures for tissue sections, plated cells, and co-localization studies can be found in the Supplementary Materials and Methods.

#### Confocal microscopy & quantification of γH2AX foci

Tissue slides or plated epithelial cells stained with γH2AX and Hoechst were imaged using a Zeiss LSM 880 confocal microscope. Confocal settings were not changed across samples within each experiment. Areas to image were first selected based on identification of regions rich in breast epithelial cells as determined by Hoechst staining prior to viewing the γH2AX channel to limit any potential bias in image selection. Images were exported to the image analysis software Imaris (Oxford Instruments) for semi-automated quantification of γH2AX foci per 100 cells. Imaris analysis settings were programmed to identify and quantify total cell number in each image and to identify number of γH2AX foci co-localizing with nuclei. All Imaris-analyzed images were visually inspected by investigators to ensure appropriate identification of γH2AX foci and exclusion of background staining. A minimum of 100 cells per case or condition were analyzed and DNA damage was reported as # of γH2AX foci per 100 cells. Any sample with less than 100 cells detected were excluded.

### Quantification of blood biomarkers

Fasting blood was collected from patients prior to surgery. Serum was separated by centrifugation, aliquoted, and stored at -80°C. Enzyme-linked immunosorbent assay was used to measure serum levels of insulin (Mercodia, Uppsala, Sweden), hsCRP, glucose, SHBG, and IL-6 (R&D Systems, Minneapolis, MN) following the manufacturer’s protocols.

### RNA-Seq studies & computational analysis

RNA-Sequencing (RNA-Seq) was conducted on samples in 4 studies including: breast tissue from *BRCA* mutation carriers, isolated breast epithelial organoids from *BRCA* mutation carriers, breast adipose tissue conditioned media (CM)-treated *BRCA*1 heterozygous MCF-10A cells, and *Brca1*+/- mouse mammary fat pads. Details on RNA extraction, sequencing methodology, and computational analyses can be found in the Supplementary Materials and Methods.

### Isolation of primary breast epithelial cells and breast explant studies

For *ex vivo* tissue explant studies and isolation of breast epithelial cells, breast tissue was obtained from women undergoing breast mammoplasty or mastectomy surgeries at Weill Cornell Medicine and MSKCC from 2017-2021. Surgical specimens were transferred from the operating room to a pathologist who evaluated the breast tissue to confirm that the tissue distributed for experimentation was normal and uninvolved with any quadrant where a tumor may have been present. The tissue was then brought to the laboratory and utilized in the experiments as described below.

#### Isolation of breast epithelial cells

Approximately 25mL of breast tissue was utilized in each organoid preparation with care taken dissect out overly fibrous areas or visible blood vessels. The tissue was finely minced and mixed with complete Ham’s F12 media (Corning #10-080-CV, supplemented with 10% FBS and 1% penicillin/streptomycin) containing a digestion mix of 10mg/mL collagenase type 1 (Sigma Aldrich #C0130) and 10µg/mL hyaluronidase (Sigma Aldrich #H3506) in a total volume of 50mL. The tissue was digested overnight on a rotator at 37°C and then centrifuged. The supernatant containing free lipid and adipocytes was discarded and the pellet was washed and reconstituted with in media followed by incubation at 4°C for 1 hour to ensure inhibition of enzyme activities. After centrifugation, the pellet was treated with red cell lysis buffer (Sigma Aldrich #11814389001), pelleted, reconstituted in media, and then ran through a 100µM filter followed by 40µM filter. Breast epithelial organoids were collected from the top of the 40µM filter in mammary epithelial cell growth medium with added supplements (PromoCell #C-21010). Isolated mammary epithelial organoids were snap frozen in liquid nitrogen for RNA extraction and RNA-sequencing or plated for *in vitro* studies.

#### Ex vivo metformin and fulvestrant explant studies

To examine the role of breast adipose tissue estrogen in mediation of DNA damage in *BRCA* mutant epithelial cells, breast explants were treated with drugs targeting estrogen signaling (fulvestrant) or production (metformin). 1 cm breast tissue explants were cut from breast tissue transferred after surgery and were plated in replicate in a 12-well dish. Metformin studies: Breast explants from n=3 subjects were cultured in complete Ham’s F12 media (10% FBS, 1% penicillin/streptomycin) supplemented with either vehicle (methanol) or metformin hydrochloride (25-100µM, Sigma #PHR1084). Fulvestrant studies: Breast explants from n=7 subjects were cultured in basal mammary epithelial cell growth media + 0.1% BSA containing either vehicle (ethanol) or 100uM fulvestrant (Sigma #I4409).

After 24 hours of treatment at 37°C in a 5% CO_2_ incubator, explants were snap frozen in liquid nitrogen and formalin fixed and paraffin embedded. Tissue sections were cut from each paraffin block for assessment of breast epithelial cell DNA damage by immunofluorescence staining.

#### Collection of breast adipose tissue conditioned media

Conditioned media (CM) was generated from breast tissue obtained from n=36 women with BMIs that range from lean to obese (20.6 – 49.1 kg/m^2^). Ten 1 cm explant pieces of breast adipose tissue were cut from each case with a focus on fatty areas containing no visible blood vessels. The pieces were weighed and placed on a 10cm dish with 10mL of basal (phenol red free, serum free, and supplement mix free) mammary epithelial cell growth media (PromoCell #C-21215) containing 0.1% BSA. The explants were incubated at 37°C for 24 hours. After incubation the breast adipose tissue CM was collected and centrifuged at 300xg. The supernatant was aliquoted and stored -80°C for use in *in vitro* treatment studies.

### *In vitro* studies in MCF-10A cells

Non-cancerous breast epithelial cell line MCF-10A carrying a *BRCA1* heterozygous mutation (185delAG/+) was purchased from Horizon Discovery and have been previously described (*66*). MCF-10A cells carrying a *BRCA2* heterozygous mutation (6174delT/+) were generated in-house using CRISPR/Cas9 gene editing (additional details provided in the Supplementary Materials and Methods). Cells were cultured in DMEM/F12 (Invitrogen #11330-032) supplemented with 5% FBS, 1% penicillin/streptomycin and the following growth factors: 20ng/mL EGF, 0.5mg/mL hydrocortisone, 100ng/mL cholera toxin, and 10µg/mL insulin (all purchased from Sigma Aldrich). Cells were serum starved for 16 hours prior to treatments.

In CM studies, CM was thawed on ice from each case and diluted to a final concentration of 25% CM. In leptin studies, cells were treated with 400ng/mL of human recombinant leptin (Sigma #L4146). In leptin neutralization studies, obese CM was pre-incubated with a leptin neutralizing antibody (Lep ab, 13.3µg/mL, Fisher Scientific #AF398) for 1 hour at 4°C and then cells were treated with lean or obese CM alone or obese CM + Lep ab. In insulin studies, cells were treated with 100nM insulin (Sigma #I1882). To block insulin signaling, cells were pre-treated with the PI3K inhibitor BKM120 (1uM, MedChemExpress #HY-70063) for 1 hour and then then treated with obese CM + BKM120. All treatments were conducted in replicates or triplicates for 24 hours unless otherwise stated. After treatment, all wells were fixed with ice cold methanol followed by γH2AX IF staining.

### *Brca1+/-* mouses studies

#### Generation of Brca1+/- mice

To determine if obesity impacts mammary gland DNA damage and tumor penetrance in the setting of a *Brca* mutation, *Brca1* heterozygous (*Brca1*+/-) mice were generated on a C57BL/6 background as described in the Supplementary Materials and Methods.

#### Diet-induced obesity & mammary gland DNA damage

At 4 weeks of age, 24 female *Brca1*+/- mice were randomized to one of two groups (n=12/gp). One group was fed 10 kcal% low fat diet (LFD, 12450Bi, Research Diets) and the second group was fed 60 kcal% high fat diet (HFD, D12492i, Research Diets) *ad libitum* for 22 weeks until sacrifice. One week prior to sacrifice all mice were fasted overnight for 12 hours and underwent glucose tolerance tests to confirm obesity-induced metabolic dysfunction as previously described. In brief, baseline glucose measurements were taken from tail vein blood drop collection using a handheld glucose meter (Bayer Contour). Mice then received an intraperitoneal injection of 1g/kg glucose and tail vein blood glucose levels were recorded at 15-30 minute intervals over 90 minutes. Following the final measurement respective experimental diets were re-started *ad libitum* for an additional week prior to sacrifice. Mice were euthanized via CO_2_ inhalation and mammary gland tissue was collected and snap frozen (inguinal fat pads) for RNA-Seq or fixed (thoracic fat pads) in 10% neutral buffered formalin overnight prior to paraffin embedding and sectioning for histological assessment of DNA damage.

#### MPA/DMBA tumor model

To investigate how obesity impacts mammary gland tumor development in *Brca1*+/- mice the same diet-induced obesity model as described above was utilized. At 4 weeks of age, 27 female *Brca1*+/- mice were randomized to one of two groups (n=13-14/gp). One group was fed LFD and the second group was fed HFD for the duration of the study. At 14 weeks of age (after 10 weeks on experimental diets) all mice were surgically implanted with a 40mg medroxyprogesterone acetate (MPA) pellet (90-day continuous release, Innovative Research of America, #NP-161) placed subcutaneously. At 15, 16, and 17 weeks of age all mice were dosed with 1mg/22g bodyweight of the carcinogen 7,12-dimethylbenz[a]anthracene (DMBA) delivered by oral gavage in corn oil once per week for 3 consecutive weeks. Mammary tumor development and growth were monitored weekly by palpating all 5 mammary gland pairs and recording tumor presence and size with caliper measurements for 28 weeks following the last dose of DMBA. Mice were euthanized at the end of the 28-week surveillance period or earlier based on ethical endpoints, including tumor burden reaching 1.5cm. Mice that did not recover from pellet implantation surgery or displayed morbidity unrelated to mammary tumors were excluded from the study.

### Quantitative steroid analysis in breast explants

Quantification of steroid levels (E2, E1, testosterone, and androstendione) in snap frozen breast adipose tissue explants treated with metformin was performed using gas chromatography-mass spectrometry (GC-MS)-based steroid profiling as previously described (*67, 68*). Detailed protocol included in the Supplementary Materials.

### Statistical analysis

To assess significant differences in baseline clinical characteristics and categorical variables the Fisher exact test was used. To test the strength of correlation between DNA damage and continuous variables, nonparametric Spearman’s rank correlation coefficient was used with two-tailed *P* value to determine significance of correlations. A multivariable linear model was used to test the association between the level of DNA damage and clinical characteristics adjusting for BMI or age. Two-tailed Mann Whitney test was performed on clinical data testing significant differences between two groups. Two-tailed student’s t-test was used *in vitro* treatment studies and in mouse studies comparing two groups. All results were performed using R (version 4.0.5) or GraphPad Prism 9. Results with a *P*-value < 0.05 were considered statistically significant.

## Supplementary Materials

Materials and Methods

**Fig. S1.** Breast adipose conditioned media from obese women stimulate more DNA damage in *BRCA1+/-* MCF-10A cells compared to conditioned media from lean women.

**Fig. S2.** *Brca1+/-* mice fed high fat diet have significantly greater accumulation of body fat compared to *Brca1+/-* mice fed low fat diet

**Fig. S3.** Generation of MCF-10A cells carrying a *BRCA*2 heterozygous mutation

**Data file S1:** Tables S1-5: Full list of differentially expressed genes in presented RNA-seq studies (multi-tab Excel file).

**Data file S2:** Original data for experiments presented

References (*69*-*80*)

## Acknowledgments

We thank Dr. David Otterburn and Dr. Leslie Cohen, Department of Surgery, and Dr. Paula Ginter, Department of Pathology, at Weill Cornell Medicine for facilitating access to reduction mammoplasty tissue. Studies were conducted with the support and facilities provided by the Microscopy and Image Analysis Core Facility, Genomics Resources Core Facility, and the Research Animal Resource Center at Weill Cornell Medicine (New York, NY). All schematics were created in BioRender.

## Author contributions

Conceptualization: KAB, NMI

Methodology: KAB, PB, NMI, HZ, DJB, MHO, QS, RB, MF, LED, DDG, XKZ

Investigation: PB, HZ, KMC, DJB, CL, CL, PP, MA, MF, SO, MP, BH, MKF

Formal analysis: KAB, PB, QS, OS, RB, XKZ

Resources: KAB, MHC, OE, AML, LHE, MM, JAS, LCC

Funding acquisition: KAB, PB

Project administration: KAB Supervision: KAB

Writing – original draft: PB, KAB

Writing – review & editing: All authors

