## Supplementary Materials & Methods for "Obesity promotes breast epithelium DNA damage in BRCA mutation carriers"

**Supplementary Materials and Methods**

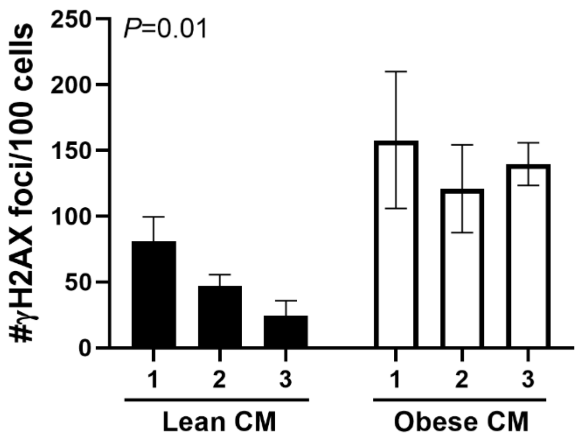


**Fig. S1**

**Fig. S1. Breast adipose conditioned media from obese women stimulate more DNA damage in *BRCA1*+/- MCF-10A cells compared to conditioned media from lean women**

*BRCA1*+/- MCF-10A cells were treated with breast adipose conditioned media (CM) collected from women identified as lean (n=3) or obese (n=3) by BMI. DNA damage was assessed by IF (#γH2AX foci/100 cells) after 24 hours treatment. *P* value represents a comparison of average DNA damage in lean CM vs obese CM treated cells determined by two-tailed student's t test. *BRCA1*+/- MCF-10A cells treated in parallel were submitted for RNA-Seq. Data is presented as mean +/- SD.

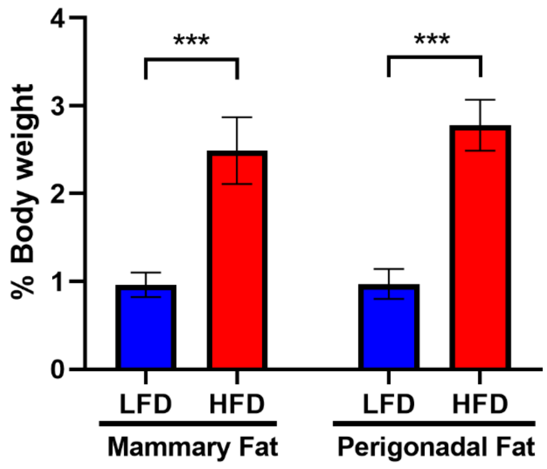


Fig. S2

**Fig. S2. *Brca1*+/- mice fed high fat diet have significantly greater accumulation of body fat compared to *Brca1*+/- mice fed low fat diet**

Wet weight of inguinal mammary fat pads (subcutaneous fat) and perigonadal fat (visceral fat) at the time of sacrifice, expressed as % of whole-body weight in female *Brca1*+/- mice fed low fat diet (LFD) or high fat diet (HFD) for 22 weeks (n=12/gp). Student's t test was used to determine significant differences between LFD and HFD mice. Data is presented as mean +/- SD. ****P*<0.001.

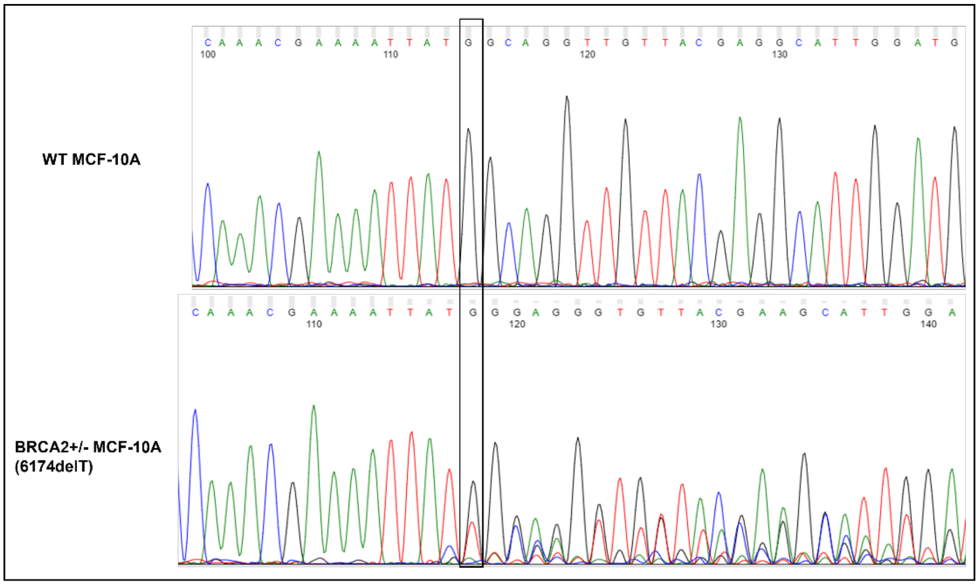


**Fig. S3**

**Fig. S3. Generation of MCF-10A cells carrying a *BRCA*2 heterozygous mutation**

CRISPR-Cas9 gene editing was utilized to generate an isogenic MCF-10A cell line carrying a heterozygous *BRCA*2 mutation (6174delT). Candidate clones were submitted for Sanger sequencing (Genewiz) and a clone exhibiting a heterozygous mutation as determined by the sequencing chromatogram shown in the figure was selected for downstream *in vitro* studies.

### Supplementary Methods

#### Immunofluorescence and immunohistochemistry staining

*Human breast, ovary, and fallopian tube tissue*: unstained sections were cut from paraffin blocks of embedded tissue or tissue microarrays. Slides were deparaffinized by immersion in xylene and rehydrated in decreasing concentrations of ethanol. Antigen retrieval was conducted by immersing the slides in citrate buffer (10mM citric acid, 0.05% Tween 20, pH 6.0), heating in a microwave, and cooling for 30 minutes at room temp. Slides were then washed, blocked for 1 hour with CAS blocking reagent, and incubated with primary γH2AX antibody overnight at 4°C. Slides were then washed and incubated with Alexa Fluor 546 secondary antibody and Hoechst stain for 2 hours. Following this incubation period, all slides were washed and mounted with ProLong Gold until imaging by confocal microscopy.

*Mouse mammary gland*: Similar protocol as described above was conducted with sections from paraffin embedded mouse mammary gland tissue, however following antigen retrieval, staining procedures were conducted in accordance with the manufacturer's protocol for the M.O.M. immunodetection kit with some modifications. In brief, slides were incubated in M.O.M. Mouse IgG blocking reagent for 1 hour, washed, and incubated with M.O.M. diluent for 5 min followed by incubation with primary γH2AX antibody in M.O.M diluent overnight at 4°C. Slides were washed, incubated with secondary antibody and Hoechst stain in M.O.M diluent for 2 hours, washed and mounted.

*Plated breast epithelial cells*: Primary breast epithelial cells or *BRCA1* and *BRCA2* heterozygous MCF-10A cells were fixed in ice cold methanol for 15 mins at -20°C, washed and blocked with CAS block reagent for 1 hour followed by overnight incubation with primary γH2AX antibody at 4°C. The next day wells were washed, incubated with secondary antibody and Hoechst for 2 hours and imaged by confocal microscopy.

*ERα IHC and IF colocalization of ERα & γH2AX*: IHC was conducted on breast tissue sections from *BRCA1* and *BRCA2* mutation carriers to detect presence of ERα positive epithelial cells. Slides were deparaffinized in xylene and re-hydrated followed by 1 hour antigen retrieval (citrate buffer 0.01 M, pH 6.0) at 95°C and 30 minutes cooling at room temperate. Slides were rinsed and then incubated for 5 minutes with 3% $H_20_2$ to inhibit endogenous peroxidase activity.

Vectastain ABC-HRP (Peroxidase, Anti-Mouse IgG) immunodetection kit (Vector Labs #PK-4002) was then used following the manufacturer's protocol. In brief, after 30-minute incubation in blocking serum, primary ERα antibody (Leica Biosystems #ER-6F11-L-F, 1:50 dilution) was added for 1 hour at room temperature followed by biotinylated secondary antibody incubation for 30 minutes. Slides were washed, developed with DAB substrate kit (Vector Labs #SK-4100) and counterstained with hematoxylin for visualization of ERα positive epithelial cells. In γH2AX co-localization studies by IF, a similar protocol through antigen retrieval was followed as described above. Slides were then blocked for 1 hour with CAS block and incubated with primary ERα and γH2AX (NB100-384, 1:300) antibodies overnight at 4°C. Slides were washed and incubated with anti-Mouse Alexa Fluor 546 and anti-Rabbit Alexa Fluor 488 secondary antibodies with Hoechst.

**RNA-Seq studies & computational analysis:**

Total RNA was extracted from samples in all studies by Trizol lysis and RNeasy Mini Kit (Qiagen) following the manufacturer's protocols. Samples were submitted for RNA-Seq at the Genomics Resources Core Facility (GRCF, Weill Cornell Medicine). Additional details for each study are described below:

*Human breast tissue*: To assess differences in breast tissue gene expression in overweight/obese (n=64) vs lean (n=67) *BRCA* mutation carriers, snap frozen non-tumorous breast tissue was obtained from women who had previously undergone prophylactic or therapeutic mastectomy surgery at MSKCC. In the therapeutic mastectomy cases, only normal breast tissue from quadrant free of tumor or contralateral breast, as determined by a pathologist, was collected and snap frozen. Sequencing libraries were constructed at the GRCF following the

Illumina TrueSeq Stranded Total RNA library preparation protocol with ribosomal RNA depletion. The libraries were sequenced with paired-end 51 bp on the Illumina HiSeq4000 sequencing platform. Raw sequenced reads were pseudoaligned to the human reference genome (UCSC hg19) using the RNA-Seq quantification program Kallisto as previously described (*69*) and transcript abundance was quantified to obtain and raw counts. Normalization of the RNAseq raw read counts and pairwise statistical analysis were performed using the DESeq2 package (version 1.30.1)(*70*). Differentially expressed genes (DEG) were determined by comparing the overweight/obese cases relative to lean cases or DNA damage Q4 relative to Q1. The tissue RNA-Seq heatmap plot (**Fig. 2A**) was generated with R pheatmap package, using the DESeq2 normalized CPM values.

DEGs with Log2FC of >0.3 and <-0.3 and *P* value<0.05 were uploaded to Ingenuity Pathway Analysis (IPA, Qiagen) for data visualization and analysis. The estrogen metabolism heatmap (**Fig. 2C**) was generated by standardizing values relative to data across cases for each gene returning a normalized value from a distribution characterized by the mean and standard deviation of each gene.

*Breast epithelial organoids*: To assess differences in breast epithelial cell gene expression in overweight/obese (n=9) vs lean (n=10) *BRCA* mutation carriers, breast epithelial organoids were isolated from patients as described in the main text Materials & Methods section. Sequencing libraries were constructed at the GRCF following the Illumina TruSeq Stranded mRNA library preparation protocol (Poly-A selection and Stranded RNA-Seq). The libraries were sequenced with paired-end 50 bp on the Illumina HiSeq4000 sequencing platform. All reads were independently aligned with STAR_2.4.0f1 (*71*) for sequence alignment against the human genome sequence build hg19, downloaded via the UCSC genome browser and

SAMTOOLS v0.1.19 (*72*) for sorting and indexing reads. Normalization of the RNA-Seq raw read counts, and pairwise statistical analysis were performed as described above. DEGs were determined by comparing the overweight/obese cases relative to lean cases. All DEGs with *P* value <0.05 were uploaded to IPA for data visualization and analysis.

*MCF-10A cells*: To assess gene expression changes associated with treatment with obese breast adipose CM, MCF-10A cells carrying a heterozygous *BRCA1* mutation were treated with lean or obese breast adipose CM (n=3/gp). RNA was extracted after 24-hour treatment for RNA-Seq. Sequencing libraries were constructed at the GRCF following the Illumina TruSeq Stranded mRNA library preparation protocol (Poly-A selection and Stranded RNA-Seq). The libraries were sequenced with paired-end 50 bp on the Illumina HiSeq4000 sequencing platform. All reads were independently aligned with STAR_2.4.0f1 (*71*) for sequence alignment against the human genome sequence build hg19, downloaded via the UCSC genome browser and SAMTOOLS v0.1.19 (*72*) for sorting and indexing reads. Cufflinks (2.0.2) (*73*) was used to estimate the expression values (FPKMS), and GENCODE v19 (*74*) GTF file for annotation. The gene counts from htseq-count and DESeq2 Bioconductor package (*70*) were used to identify DEGs. All DEGs with *P* value <0.05 were uploaded to IPA for data visualization and analysis.

*Mouse mammary fat pads*: To assess differences in mammary fat pad gene expression in association with obesity, mammary fat pads were harvested from female *Brca1*+/- mice fed high fat diet-fed or low fat diet-fed (n=6/gp) followed by RNA extraction for RNA-Seq. Sequencing libraries were constructed at the GRCF following the Illumina TruSeq Stranded mRNA library preparation protocol (Poly-A selection and Stranded RNA-Seq). The libraries were sequenced with paired-end 50 bp on the Illumina NovaSeq 6000 sequencing platform. The raw sequencing reads in were processed through bcl2fastq 2.19 (Illumina) for FASTQ conversion and demultiplexing. After trimming the adaptors with cutadapt (version1.18), RNA reads were aligned and mapped to the GRCm38 mouse reference genome by STAR (Version2.5.2) (*71*), and transcriptome reconstruction was performed by Cufflinks (Version 2.1.1). The abundance of transcripts was measured with Cufflinks in Fragments Per Kilobase of exon model per Million mapped reads (FPKM). Normalization of the RNA-Seq raw read counts and pairwise statistical analysis were performed using the DESeq2 package (version 1.30.1) (*70*). DEGs were determined by comparing the high fat diet group relative to the low fat diet group.

Heatmaps showing comparisons between RNA-seq data from mouse and human mammary tissue were generated using IPA. Briefly, mouse raw read counts were standardized relative to data across all mouse samples for each gene returning a normalized value from a distribution characterized by the mean and standard deviation of each gene. Top canonical pathways affected based on DEGs from human overweight/obese cases relative to lean cases were then compared to standardized values from mouse samples.

**Generation of Brca1+/- mice**

We generated global *Brca1* heterozygous mice (*Brca1+/-*) by crossing *Brca1*$^{flox\ 5\text{-}13/flox\ 5\text{-}13}$ (*75, 76*) with CMV-Cre mice (JAX strain #006054, (*77*)) to produce global *Brca1*$^{\mathrm{D5\text{-}13/+}}$ mice. Genotyping was performed via PCR with primers as previously described (*75*). No offspring were found to be homozygous knockout for *Brca1* suggesting that complete loss of *Brca1* is embryonically lethal. These mice were backcrossed to a C57Bl/6 strain over 20 times. Genetic testing of mice following backcrossing was performed at Charles River Genetic Testing Services and demonstrated 98.6% fidelity with C57Bl/6 inbred strains.

### Generation of *BRCA2* heterozygous MCF-10A cell line

We used CRISPR-Cas9 gene editing to generate an isogenic MCF-10A cell line carrying a clinically-relevant heterozygous *BRCA*2 mutation (6174delT). The forward (A) and reverse (B) sgRNA cloning primers were as follows:

A: CACCGGCCAAACGAAAATTATGGC

B: AAACGCCATAATTTTCGTTTGGCC

These primers were annealed and cloned following standard procedures using the BsmBI/EcoRI site of the *pLenti-U6-sgRNA-tdTomato-P2A-Blas* (LRT2B) vector. WT MCF-10A cells stably expressing an optimized *pLenti-Cas9-P2A-Puro* lentiviral vector were transduced with sgRNAs and underwent Blasticidin selection followed by isolation of single clones using the limiting dilution assays, as described previously (*78, 79*). Candidate colonies expressing TdTomato were submitted for Sanger sequencing (Genewiz) to identify clones exhibiting a heterozygous *BRCA*2 mutation (**Fig. S3**). Analysis of sequencing data showed insertion of a thymine (T) nucleotide 5670bp from the start of the open reading frame (ATG) that led to an early stop codon at amino acid 1889. This mutation was maintained upon re-sequencing after multiple passages. The clonal population was expanded and utilized for downstream *in vitro* studies.

### Quantitative steroid analysis in breast explants

Snap frozen breast tissue explants were lyophilized and homogenized and 10 mg of individual samples were used in each assay. After spiking with 20 µL of an internal standard mixture

(2,2,4,6,6,21,21,21-$d_8$-17α-hydroxyprogesterone, 1 μg/mL; 2,2,3,4,4,6-$d_6$-dehydroepiandrosterone and 2,2,4,6,6,17α,21,21,21-$d_9$-progesterone, 0.5 μg/mL; 16,16,17-$d_3$-testosterone and 9,11,12,12-$d_4$-cortisol, 0.25 μg/mL; 2,4,16,16-$d_4$-17β-estradiol, 0.2 μg/mL) and mixed with 1 mL of 0.2 M phosphate buffer (pH 7.2), the sample was pulverized using a TissueLyser (Qiagen, Hilden, Germany) at 25 Hz for 10 min with three zirconia beads (3.0 mm I.D., Toray Industries, Tokyo, Japan) and centrifuged at 12,000 rpm for 10 min twice. The combined supernatant was loaded onto a preconditioned solid-phase extraction (SPE) cartridge (Oasis HLB; 3 mL, 60 mg; Wasters, Milford, MA, USA), the SPE cartridge was washed with 1 mL of water twice and eluted with 2 mL of methanol and 2 mL of 90% methanol. The combined eluate was evaporated under a nitrogen stream at 40°C. The sample was dissolved in 1 mL of 0.2 M acetate buffer (pH 5.2) and 100 µL of 0.2% ascorbic acid solution. It was then extracted with 2.5 mL of methyl *tert*-butyl ether twice. The organic solvent was evaporated under a nitrogen stream and further dried in a vacuum desiccator with $P_2O_5$/KOH for at least 30 min. Finally, the dried residue was derivatized with 50 µL of *N*-methyl-*N*-trifluorotrimethylsilylacetamide/ammonium iodide/dithioerythritol (500:4:2, v/w/w) at 60°C for 20 min. Then 2 µL of the final mixture was injected into the GC-MS system.

In samples with no quantifiable estrogens, the estrogen assay with improved analytical detectability (*80*) was performed from an additional 10 mg of homogenized breast tissues. For the calibration sets, steroid-free breast tissue was freshly prepared 1 day before the experiment. Human breast samples (50 mg) were pulverized in 1 mL methanol/chloroform (1:1, v/v) with 4 zirconia beads at 25 Hz for 5 min followed by centrifugation twice at 12,000 rpm for 3 min, and then supernatants were discarded. The remaining tissue sample was washed with l mL of chloroform/0.6 M methanolic HCl (1:1, v/v), sonicated for 5 min, and centrifuged at 12,000 rpm for 3 min three times. All supernatants were also discarded. To eliminate residual methanolic HCl, 1 mL of 20% ethanol was added for washing five times. Tissue samples were then frozen at -80 °C until used. No steroid was detectable in GC-MS chromatogram.
